# Chromosome-level Atlantic switchgrass assemblies reveal a supernumerary B-Chromosome with transcriptional and evolutionary impacts

**DOI:** 10.64898/2026.10.17.759809

**Authors:** Ahmed F. Elfarargi, Li Zhang, Adam L. Healey, Jason E. Bonnette, Avinash Sreedasyam, Usha Bishnoi Saran, Jerry Jenkins, David B. Lowry, John T. Lovell, Kerrie Barry, LoriBeth Boston, Jie Guo, Jayson Talag, Sumaira Zaman, Jane Grimwood, Jeremy Schmutz, Thomas E. Juenger

## Abstract

Supernumerary B-Chromosomes are extra genetic elements that frequently deviate from Mendelian inheritance to ensure their own survival. Their evolutionary origins and biological impacts within natural plant populations remain largely unexplored. Switchgrass (*Panicum virgatum*) is a model for studies of local adaptation, but reliance on a single reference genome has limited the detection of lineage-specific genomic content. We generated chromosome-level genome assemblies from the diverged Atlantic switchgrass subpopulation and discovered a previously undetected genomic element: a ∼34 Mb supernumerary B-Chromosome, validated by Hi-C contact mapping and long-read depth, which indicates two to four B copies per nucleus. The B-Chromosome has a bipartite architecture including a gene-rich, single-copy proximal region (∼8 Mb) and a repeat-rich, gene-poor distal region (∼27 Mb) dominated by LTR/Ty3 (Gypsy) elements and punctuated by Mutator TIR peaks. High-resolution mapping reveals a highly rearranged chimera of fragments from multiple autosomes, biased toward the N-subgenome. Intact LTR retrotransposons on the B-Chromosome are younger than those on the A-Chromosomes, especially in its distal region, consistent with a B-Chromosome that arose and expanded within the Atlantic lineage. Population screening and phylogenetic analysis indicate that the B-Chromosome is largely restricted to the Atlantic subpopulation and likely has a single origin. In a typical family, 70% of open-pollinated progeny of B-positive maternal plants were B-positive, with substantial variation among families. Finally, B-Chromosome presence is associated with expression differences in A-Chromosome genes that differ between genetic backgrounds. Together, our results show how accessory chromosomes contribute to the evolutionary and genomic diversity of natural plant populations.

## Introduction

Switchgrass (*Panicum virgatum*), a perennial C4 grass, is a species of significant dual importance, serving as both a foundational component of North American tallgrass prairie ecosystems and a leading lignocellulosic feedstock for sustainable biofuel production (Sanderson et al. 1996; Schmer et al. 2008). Its evolutionary success is exemplified by its vast distribution across the steep climatic gradients of North America, establishing it as a model system for dissecting the genetic architecture of local adaptation (McMillan 1964; Casler et al. 2004; Casler et al. 2007; Lowry et al. 2019; Lovell et al. 2021). The species also exhibits substantial genomic plasticity: like other polyploid panicoid grasses, switchgrass occurs at multiple ploidy levels, most commonly tetraploid (2n = 4x = 36) and octoploid (2n = 8x = 72), a flexibility thought to contribute to its wide habitat range and resilience (Comai 2005; Napier et al. 2022). Within the species, population genomic studies have resolved deeply divergent genetic lineages, including distinct Midwest, Gulf, and Atlantic subpopulations (Lovell et al. 2021; Napier et al. 2022) and considerable ecotypic divergence. The Atlantic subpopulation, occupying a narrow coastal corridor that nonetheless spans a broad latitudinal range from the southeastern to the northeastern United States, has not only evolved in response to strong local environmental pressures but has also been shaped by adaptive introgression from the pre-adapted northern Midwest gene pool, which facilitated its post-glacial colonization of northern habitats (Lovell et al. 2021). While this genetic diversity is a crucial resource for crop improvement and understanding plant adaptation, it also presents significant challenges for genomic analysis and for understanding the evolution of the switchgrass ecotypes.

To date, genomic studies in switchgrass, including large-scale analyses of climate adaptation, have predominantly relied on a single high-quality reference genome derived from a single genotype (AP13) of the lowland cultivar “Alamo”, which belongs to the southern Gulf subpopulation (Lovell et al. 2021; Napier et al. 2022). While this reference has been invaluable, it limits the resolution of lineage-specific variation across the species’ divergent gene pools, including the Atlantic and Midwest subpopulations, owing to the substantial evolutionary divergence between them, estimated to be over half a million years (Lovell et al. 2021). This limited resolution can mask lineage-specific structural variants, complicate the analysis of gene expression, and obscure unique genomic features that may be critical to local adaptation in distinct subpopulations. The absence of high-quality, chromosome-level reference genomes presenting the scope of natural diversity has therefore represented a critical gap in our ability to fully explore the unique evolutionary history and adaptive potential of switchgrass.

Beyond changes in whole-genome ploidy, a distinct and far less understood class of genomic elements, supernumerary B-Chromosomes, can also restructure plant genomes. These accessory chromosomes are found across thousands of eukaryotic species and can exhibit complex evolutionary dynamics, often maintained in populations despite lacking essential housekeeping genes (Jones 1995; Camacho et al. 2000; Ferree et al. 2024). However, our current mechanistic understanding of B-Chromosomes is heavily biased toward a few major crop species, such as maize (*Zea mays*) and rye (*Secale cereale*), where the availability of reference assemblies has enabled detailed molecular interrogation (Martis et al. 2012; Birchler and Yang 2021; Blavet et al. 2021). In these model systems, B-Chromosomes are now known to be transcriptionally active, capable of modulating the expression of the A-Chromosome complement, and equipped with sophisticated non-Mendelian accumulation mechanisms, such as nondisjunction during pollen mitosis, to ensure their persistence (Parker 1976; Jones 1995; Banaei-Moghaddam et al. 2012; Huang et al. 2016; Valente et al. 2017; Blavet et al. 2021; Johnson Pokorná and Reifová 2021; Chen et al. 2022; Ferree et al. 2024). In grasses particularly, B-Chromosomes have been documented for decades through cytogenetic studies (D’Ambrosio et al. 2017), while their genomic architecture and functional impact in natural populations remain largely unknown. Consequently, the potential for such a dynamic genomic element to exist and function within the genome of switchgrass has, until now, been unexplored.

Here, we present two high-quality, chromosome-level genome assemblies from the Atlantic switchgrass subpopulation, providing a vital resource for lineage-specific genomic research. Using these new references, we report the discovery and complete assembly of a supernumerary B-Chromosome, validate it with chromatin-contact data and long-read sequencing depth, and estimate its copy number. We also provide a high-resolution map of the five genetic subgroups that define the Atlantic subpopulation. We characterized the B-Chromosome’s genomic landscape, revealing its gene and repeat content in contrast to the core A-Chromosome complement. Through phylogenetic and syntenic analyses, we tested competing hypotheses for its evolutionary origin, finding evidence consistent with a single origin within the Atlantic lineage. We also screened the species-wide diversity panel for the B and measured how often the progeny of B-positive plants carry it. Finally, by integrating transcriptomic data, we investigated the B-Chromosome’s transcriptional footprint to assess its potential functional role within the Atlantic subpopulation.

## Results

### Chromosome-level Atlantic genome assemblies reveal a supernumerary B-Chromosome

To provide a lineage-specific genomic resource for the diverged Atlantic subpopulation, we generated a new, high-quality, chromosome-level reference genome from the Atlantic genotype J621C. We also assembled the genome of a second Atlantic genotype, J586A, which was collected from a geographically and genetically distinct part of the Atlantic range (a southern genotype, versus the northern/Chesapeake-associated J621C). Sequencing two distinct Atlantic genotypes allowed us to better capture the genomic content and divergence of this lineage relative to the existing reference and, as described below, provided independent replication of a novel genomic element. By comparing these resources with the existing Gulf reference (AP13), we characterize genomic features of the Atlantic subpopulation that a Gulf-derived reference cannot capture (**Table 1**; **Table S1**).

**Table 1.** Genome assembly and annotation statistics for the two new Atlantic references versus the Gulf reference AP13 v5.0 (Lovell et al. 2021). Values for the Atlantic genotypes are shown as HAP1 / HAP2.

| Genomic feature and metric | *P. virgatum* (AP13 v5.0) | *P. virgatum* (J621C) HAP1 / HAP2 | *P. virgatum* (J586A) HAP1 / HAP2 |
| --- | --- | --- | --- |
| Ecogeographic lineage | Gulf (Lowland) | Atlantic (Chesapeake; MD) | Atlantic (Southern; SC) |
| A-Chromosomes (Standard) | 18 | 18 / 18 | 18 / 18 |
| B-Chromosomes (Supernumerary) | 0 | 1 / 1 | 1 / 0 |
| B-Chromosome copies per nucleus (HiFi depth) | NA | ≈2 (1.9) | ≈4 (3.8) |
| Genome size (Mb) | 1130 | 1188.2 / 1142.3 | 1165.2 / 1115.7 |
| No. of contigs | 1090 | 50 / 83 | 47 / 42 |
| Contig N50 (Mb) | 5.5 | 51.2 / 50.1 | 51.2 / 51.8 |
| Scaffold N50 (Mb) | 64.3 | 65.7 / 68.2 | 63.8 / 63.7 |
| GC content (%) | 46.8 | 46.8 / 46.8 | 46.8 / 46.9 |
| Protein-coding genes (Total) | 80278 | 60257 / 59975 | 58019 / 57441 |
| Protein-coding genes (B-Linked) | NA | 207 / 106 | 193 / NA |
| Total repeat content (%) | 55.8 | 57.2 / 56.3 | 57.7 / 56.8 |
| B-Chromosome repeat content (%) | NA | 79.5 / 75.1 | 79.5 / NA |
| BUSCO completeness (%) | 99.4 | 99.6 / 99.7 | 99.7 / 99.6 |

While most sequences were mapped to the standard 18 chromosomes (A-Chromosomes), an additional chromosome-scale scaffold was assembled exclusively in the two Atlantic reference genomes (J621C and J586A). This additional sequence, designated Chr19, was entirely absent from the existing Gulf AP13 genome reference (**Fig. 1**), marking the discovery of a new ∼34 Mb accessory chromosome presence-absence polymorphism in switchgrass.

**Fig. 1.**
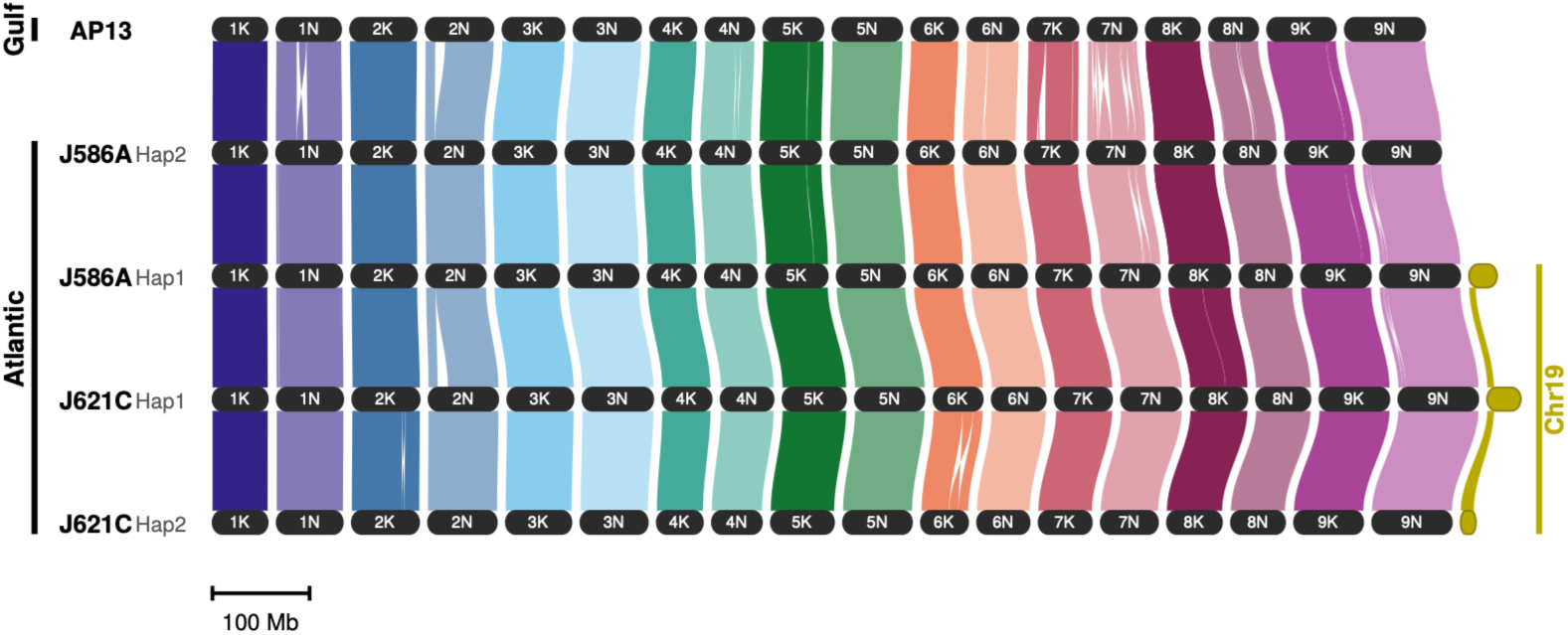
Atlantic switchgrass genome references and the discovery of a supernumerary B-Chromosome. GENESPACE synteny riparian plot of five chromosome-level assemblies: the Gulf reference AP13 (v5.1) and the haplotype-resolved Atlantic genotypes J586A (Hap2, Hap1) and J621C (Hap1, Hap2), ordered top to bottom. Braids connect syntenic gene blocks between vertically adjacent haplotypes and are colored by chromosome; crossing braids indicate inversions and translocations, and black gaps mark segments with no syntenic counterpart in the neighboring haplotype. The supernumerary B-Chromosome (Chr19) is present in J586A Hap1 and in both J621C haplotypes and has no counterpart in AP13. All chromosomes are drawn on one physical scale and separated by equal gaps, so that the length of each row reflects the size of that genome (scale bar, 100 Mb).

The independent recovery of Chr19 in two genotypes from geographically distinct regions argues strongly against an assembly artifact. We confirmed this directly with genotype-specific chromatin-contact data: Hi-C (Omni-C) contact maps show that Chr19 forms a self-contained contact domain distinct from the 18 A-Chromosomes (both the K and N subgenomes) in both J621C (**Fig. S1A**) and J586A (**Fig. S1B**), supporting its assembly as an independent chromosome rather than a mis-join or scaffolding error. Long-read sequencing depth independently supports the B as a genuine chromosome and shows that it is present in multiple copies. We aligned the PacBio HiFi reads of each genotype to a combined reference containing both haplotype assemblies and all of its assembled B copies, so that each A-Chromosome sequence represents a single copy per diploid nucleus (Methods). Normalized depth was uniform across all 36 A-Chromosome sequences of each genotype (0.97–1.05 of the single-copy depth), whereas the B was strongly over-represented (**Fig. S1C**; **Table S2**). In J586A, B3 had a median depth of 158× against a single-copy A-Chromosome depth of 41×, corresponding to 3.82 B copies per diploid nucleus (95% CI 3.72–3.95; 3.83 in uniquely mappable windows), that is, about four copies, or twice the depth of the diploid autosomes. In J621C, the part of B1 absent from B2 carried 1.89 copies (95% CI 1.83–1.95), and the region shared by B1 and B2 carried 2.03 copies once the reads divided between the two near-identical assembled copies were combined, indicating about two full-length B copies per nucleus (**Table S2**). The two genotypes thus differ in B number, a numerical polymorphism characteristic of B-Chromosomes. The recovery of the element in two independent Atlantic assemblies immediately raised the question of how prevalent it is across the broader diversity of switchgrass, which we address below with a genome-wide *k*-mer screen.

The assembled B sequences vary in size and structure: J621C carries a 34.8-Mb copy on HAP1 (B1, the reference used below) and a shorter alternate 15.7-Mb copy placed in HAP2 (B2), whereas J586A carries a single 29.6-Mb copy (B3). The three copies share substantial but incomplete syntenic and rearranged segments (49% of B1 and 99% of B2 align between B1 and B2, while 60% of B1 and 72% of B3 align between B1 and B3). Within these aligned regions, the copies are nearly identical (833 SNPs between B1 and B2, ∼0.05 per aligned kb; ∼1.4 SNPs per aligned kb between B1 and B3) (**Fig. S2**; **Table S3**). Complementary comparisons show shared sequence structure between B1 and B2 and between B1 and B3 (**Fig. S3A** and **S3B**). Self-comparisons reveal relatively low-repeat regions of approximately 8 Mb in B1 and B3 and approximately 4.1 Mb in B2, followed by extensive internal repeat matches (**Fig. S4A-C**). Furthermore, we identified an approximately 1.4 Mb insertion in B2 relative to B1. These results support structural relatedness among the assembled Bs without establishing a unique origin. HiFi read depth shows that J621C carries two full-length B copies and that B2 is shorter only because the second copy was not fully assembled: the part of B1 that is missing from B2 is present in about two copies per nucleus (**Table S2**).

The two Atlantic assemblies are highly contiguous and complete (∼100% of assembled bases integrated into chromosomes; transcript completeness of 91.5% and 90.9%), exceeding the contig contiguity of the Gulf AP13 reference while resolving a lineage that AP13 cannot represent (**Table S1**). Scaffold contiguity, base composition, repeat content and gene-space completeness are comparable between the Atlantic and Gulf references, whereas contig contiguity is about tenfold higher in the Atlantic assemblies (**Table 1**): contig N50 of ∼50-52 Mb in each Atlantic haplotype versus ∼5.5 Mb in AP13, scaffold N50 of ∼64-68 Mb versus ∼64 Mb, GC content of 46.8-46.9% in all assemblies, repeat content of 56-58% versus 56%, and 99.6-99.7% complete BUSCOs versus 99.4%. The Atlantic haplotypes carry ∼57,000-60,000 annotated protein-coding genes versus ∼80,000 in AP13, a difference that most likely reflects annotation-pipeline versions rather than biological gene loss (Brůna et al. 2026). The clearest genome-level difference between the Gulf and Atlantic references is therefore not overall size or scaffold contiguity but the presence of the accessory Chr19 in the Atlantic references and its absence from AP13 (**Table 1**).

### Genomic architecture and evolutionary age of the B-Chromosome

Detailed characterization of the B-Chromosome’s genomic content revealed that it is substantially smaller than the standard switchgrass autosomes (approximately 34 Mb compared with 56-82 Mb for the A-Chromosomes) and carries 207 predicted protein-coding genes on the complete B1 assembly, with 106 genes on the shorter B2 copy and 193 on B3 (**Table S4**; **Table 1**). Repeat annotation covers ∼80% of B1 and B3 (75% of B2), compared with 56-57% of the A-Chromosomes (**Table 1**). We annotated repeats with the Extensive de-novo TE Annotator (EDTA) and summarized the annotated fraction of overlapping 500-kb windows along B1. This analysis revealed a stark bipartite landscape (**Fig. 2A**). The first approximately 8 Mb is relatively gene-rich, whereas the remaining approximately 27 Mb is repeat-rich and gene-poor. In contrast to the A-Chromosomes, each of which carries a single, highly localized pericentromeric block dominated by Mutator TIR elements (**Fig. S5**), the distal region of the B-Chromosome is enriched in repetitive elements along its whole length. A comparison of the composition of the annotated sequence of the A- and B-Chromosomes (**Fig. 2B**) highlights an expansion of LTR/Ty3 (Gypsy) retrotransposons on Chr19, which make up 66% of its annotated sequence versus 28-40% on the A-Chromosomes, whereas genes make up only 3% (17-35% on the A-Chromosomes) (**Table S5**). Most notably, the distal part of the chromosome is punctuated by recurrent, localized high-density spikes of Mutator TIR transposable elements (**Fig. 2A**). A candidate 166-bp centromeric repeat identified on the A-Chromosomes also marks two intervals on B1, at approximately 2.87–3.04 Mb and 25.7–25.85 Mb (**Table S6**); the latter overlaps a Mutator TIR peak. Because plant centromeres are specified epigenetically by the histone variant CENH3 rather than by DNA sequence (Henikoff et al. 2001; Talbert et al. 2002; Fu et al. 2013; Su et al. 2018), these intervals are candidate positions only; which of them, if either, forms the functional centromere of the B remains to be tested by CENH3 chromatin immunoprecipitation or immunostaining, the approaches that located the functional centromeres of rice and maize chromosomes and of the maize and rye B-Chromosomes (Nagaki et al. 2004; Banaei-Moghaddam et al. 2012; Gent et al. 2017; Liu et al. 2025).

**Fig. 2.**
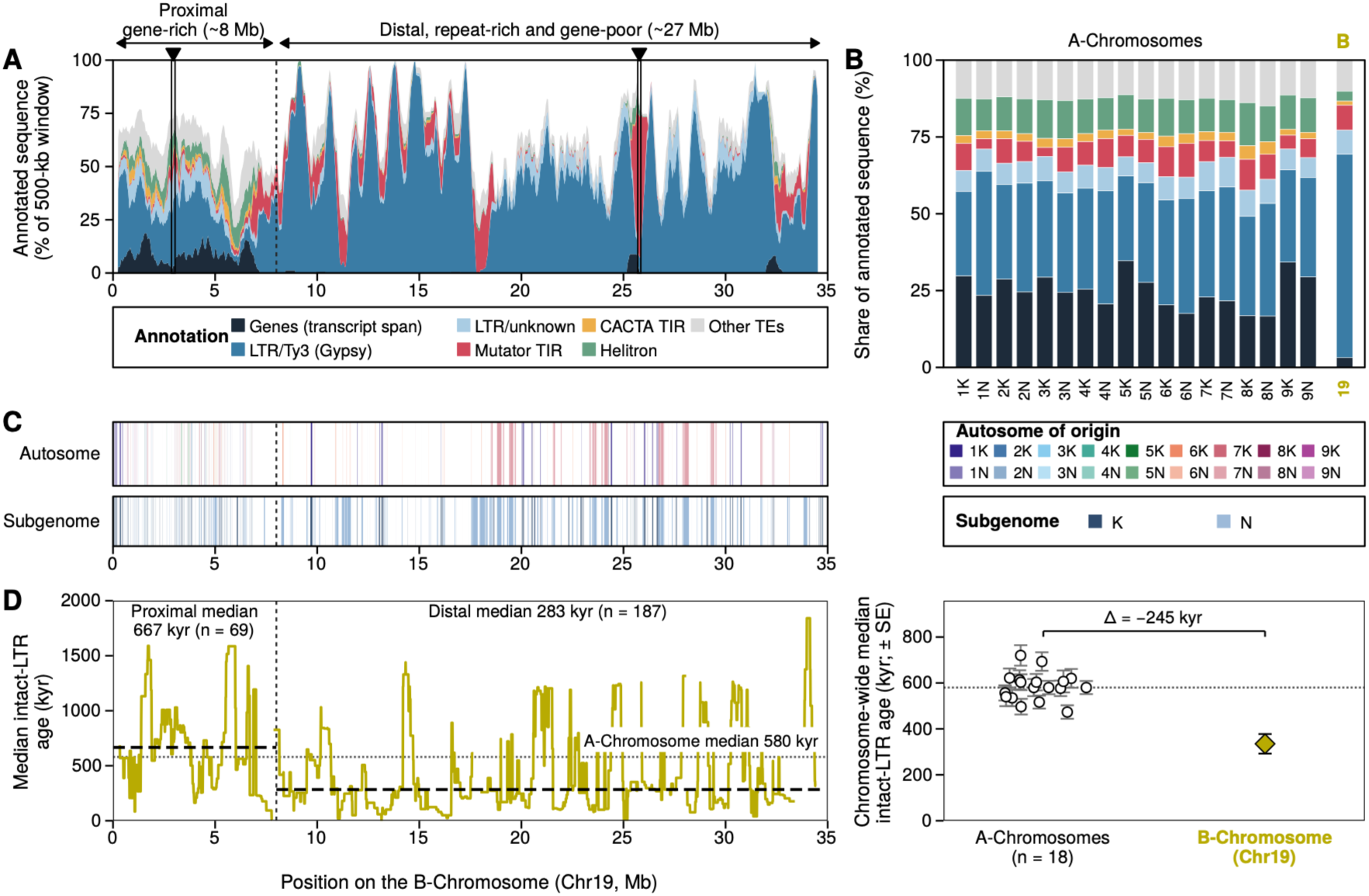
Genomic architecture, sequence origin and evolutionary age of the Atlantic B-Chromosome. (**A**) Annotation landscape of the B-Chromosome (Chr19) of J621C HAP1: percentage of each 500-kb window (moved in 50-kb steps) annotated as genes (primary-transcript spans) or as each class of transposable element, showing the gene-rich proximal region (∼8 Mb) and the repeat-rich, gene-poor distal region (∼27 Mb); boxes and arrowheads mark the two candidate centromeric-repeat intervals (**Table S6**). Each base is assigned to one class, so overlapping annotations are counted once; other TEs, LTR/Ty1 (Copia), hAT, PIF/Harbinger and Tc1/Mariner elements. The equivalent plots for the 18 A-Chromosomes are shown in **Fig. S5**. (**B**) Share of the annotated sequence of each chromosome made up by each class, 18 A-Chromosomes versus the B (Chr19), computed from the same per-base assignment as A (**Table S5**). (**C**) Origin of the B-Chromosome inferred from single-copy B-specific 80-mers realigned to the A-Chromosomes: upper track, autosome of origin (colors as in Fig. 1); lower track, subgenome of origin. Each bar is a run of at least 200 consecutive markers with the same best A-Chromosome hit (**Table S7**). (**D**) Left: median intact LTR insertion age (thousand years, kyr) along the B-Chromosome in 500 kb windows stepped by 10 kb. Dashed horizontal lines mark the median ages of the proximal (<8 Mb; 667 kyr) and distal (≥8 Mb; 283 kyr) regions, and the dotted line marks the A-Chromosome median (580 kyr). Right: chromosome-wide median intact LTR age (± bootstrap standard error) of each of the 18 A-Chromosomes (open circles) and of the B (Chr19; diamond); Δ, difference between the B median and the pooled A-Chromosome median. Panels A, C and D (left) share one Chr19 position axis, and the dashed vertical line marks the proximal/distal boundary at 8 Mb. A-Chromosome age profiles are in **Fig. S7**; values in **Table S8**.

To determine the physical sequence origins of the B-Chromosome, we used a *k*-mer-based approach to map B-Chromosome-derived sequences back to the A-Chromosomes. Single-copy B-specific 80-mers were realigned to the A-Chromosomes and their best hits projected back onto Chr19, painting the B by subgenome and by autosome of origin (**Fig. 2C**). Of the 7.87 Mb of B sequence that could be assigned confidently to one subgenome, 6.02 Mb (76.5%) derives from the N subgenome. At the level of individual chromosomes, 3.74 Mb could be traced to a specific donor autosome, and three autosomes account for 88% of this sequence: Chr07N (58%), Chr01K (19%) and Chr06N (10%) (**Table S7**). The B is therefore a highly rearranged chimera built predominantly, but not exclusively, from N-subgenome material. Gene synteny analysis independently shows that the B-Chromosome is a mosaic of conserved gene blocks derived from multiple chromosomes, including Chr01, Chr02, Chr07, and Chr09, from both subgenomes (K and N) (**Fig. S6**). The two approaches agree on the leading donors (Chr07 and Chr01) and differ only in the weight given to low-coverage contributors, as expected given that gene synteny samples the gene-bearing proximal region while the *k*-mer painting samples all single-copy sequence.

To investigate the evolutionary timeline of the B-Chromosome, we estimated its age using Long Terminal Repeat (LTR) retrotransposon divergence. Intact LTR elements extracted from the EDTA annotation were dated from the divergence of their two terminal repeats as T = d/2r, where d is the divergence between the two repeats and r = 1.5 × 10⁻⁸ substitutions per site per year (Koch et al. 2000). The median age of intact LTR elements on the B-Chromosome (335 thousand years [kyr]) was lower than on any of the 18 A-Chromosomes (per-chromosome medians 473-720 kyr; pooled median 580 kyr) (**Fig. 2D**; **Fig. S7**; **Table S8**). Within the B, intact LTRs in the gene-rich proximal ∼8 Mb had a higher median age (667 kyr) than those in the repeat-rich distal region (283 kyr) (**Fig. 2D**). Together with the gene and repeat densities (**Fig. 2A**), this suggests that the proximal region retains sequence derived from A-Chromosomes, whereas the distal region was built up more recently, largely by the accumulation of LTR/Ty3 retrotransposons. Because insertion ages date the accumulation of repeats rather than the formation of the B itself, and depend on the assumed substitution rate, they bound rather than establish the age of the element. The presence of genetic material from both subgenomes on the B most plausibly places its formation after the K and N subgenomes were united in the allotetraploid ancestor (≤4.6 MYA; Lovell et al. 2021). An LTR complement younger than that of every A-Chromosome, together with the near-restriction of the B to the Atlantic subpopulation (see below), is most consistent with a B-Chromosome that arose, and continues to accumulate repeats, within the Atlantic lineage.

### Evolutionary history, spatial distribution, and frequency of B-positive progeny

To investigate the genetic landscape of the Atlantic subpopulation at high resolution, we re-analyzed whole-genome resequencing data from 280 Atlantic genotypes sampled from natural populations and previously assigned to this subpopulation in (Lovell et al. 2021); breeding selections and cultivars were excluded (Methods; **Table S9**). Raw sequencing reads were mapped to our new chromosome-level J621C assembly to generate a set of variant calls specific to the Atlantic gene pool. Using this lineage-specific dataset, we conducted a model-based clustering (ADMIXTURE) analysis (**Fig. S8**) and a broad-scale characterization using principal component analysis (PCA) (**Fig. 3A**). The cross-validation plot from ADMIXTURE clustering shows that the optimal number of genetic clusters is five (K = 5) (**Fig. S8B**; **Table S10**). The map (**Fig. S8A**) shows how these five genetic clusters are distributed into ecogeographic regions: the Northern Coastal Plains, Mid-Atlantic Coastal Plains, Chesapeake Coastal Core, Atlantic-Midwest Admixture Group, and Southern Coastal Plains. We also confirmed this genetic structure with the PCA, which separates the same five subgroups in genetic space; the Northern Coastal Plains, Chesapeake Coastal Core, and Southern Coastal Plains each form a distinct cluster (**Fig. 3A**; **Table S9**). In contrast, individuals from the Mid-Atlantic Coastal Plains and Atlantic-Midwest Admixture Group did not form discrete clusters but instead occupied intermediate positions in PCA space, bridging the distinct Northern, Chesapeake, and Southern groups. Notably, the intermediate position of the Atlantic-Midwest Admixture Group aligns with previous genomic findings of adaptive introgression from the Midwest lineage into the Atlantic subpopulation (Lovell et al. 2021).

**Fig. 3.**
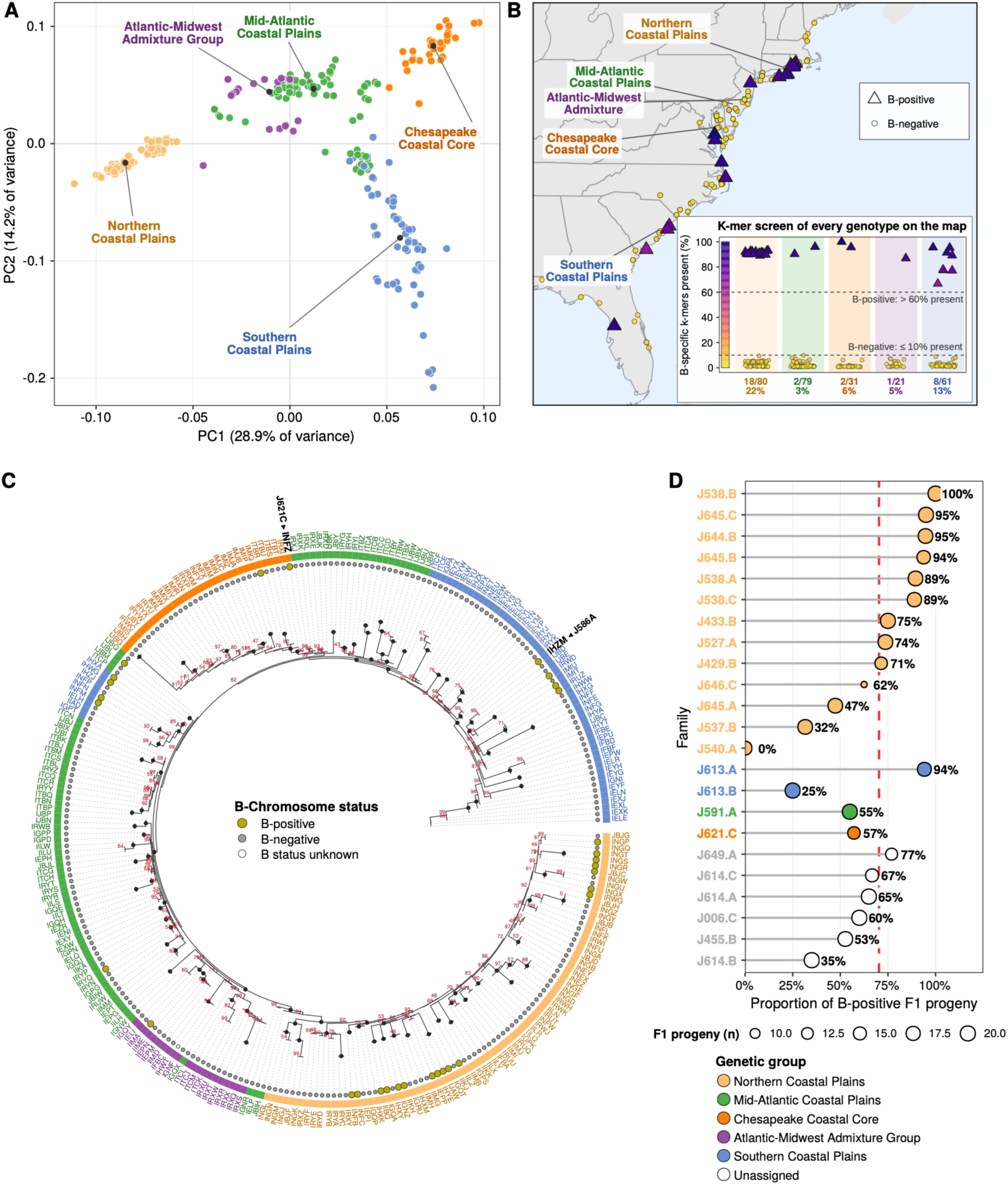
B-Chromosome distribution, origin, and frequency in progeny. (A) Principal Component Analysis (PCA) based on genome-wide SNPs, illustrating the genetic differentiation among the five Atlantic switchgrass groups. Individuals are plotted along the first two principal components (PC1 and PC2) and colored according to their primary genetic group as determined by ADMIXTURE analysis (**Fig. S8**; key below D, which also applies to A and C). (**B**) Geographic distribution of B-positive genotypes (triangles) and B-negative genotypes (circles) along the Atlantic coast, identified by the k-mer screen; every symbol is filled by the percentage of B-specific k-mers present in its sequencing reads (color scale along the y axis of the inset), and the genetic groups are labeled on the map. The inset plots every mapped genotype with the same symbols and colors against the percentage of B-specific k-mers present, by genetic group, with the classification thresholds (B-positive, > 60% of B-specific k-mers present; B-negative, ≤ 10%) and the number and percentage of B-positive genotypes in each group. (**C**) Coalescent-aware phylogenetic tree of the Atlantic subpopulation inferred using CASTER-site. Black dots mark nodes with 100% local bootstrap support; support values below 100% are given as numbers. The outer ring colors correspond to the five genetic clusters defined in (**A**). Gold points beside the tips indicate B-positive genotypes (grey, B-negative; open circle, B status unknown), highlighting the distribution of the B-Chromosome across distinct genetic groups. The two assembled genotypes, J621C and J586A, are labeled. (**D**) Proportion of B-positive F1 progeny in 23 open-pollinated families of B-positive maternal plants. Point sizes correspond to the number of F1 progeny screened per maternal line. Colors give the genetic group of the maternal plant (open circles, unassigned). The vertical red dashed line marks the B-positive frequency for a typical family estimated with a Generalized Linear Mixed Model (GLMM; 70.3%, 95% CI 56-81%). The among-family variance (random intercept variance = 1.91 on the logit scale) indicates substantial variation among families.

Having established the population structure of the Atlantic gene pool, we next evaluated the prevalence and evolutionary history of the B-Chromosome. We first screened the switchgrass diversity panel of Lovell et al. (2021) (696 genotypes with sequencing libraries, from the Gulf, Midwest, and Atlantic subpopulations) for single-copy 80-mers of the J621C B-Chromosome that are absent from the A-Chromosomes of eight switchgrass assemblies (Methods; **Fig. 3B**; **Table S11**). The B-Chromosome was predominantly associated with the Atlantic subpopulation: it was carried by 38 of the 288 Atlantic genotypes of the panel with a confident B-status call (13.2%) and by 31 of the 272 such genotypes in our 280-genotype Atlantic set (11.4%). In contrast, it was detected in only 2% (3 of 153) of the panel’s Gulf genotypes and in none of its 197 Midwest genotypes (**Table S12**), suggesting a unique evolutionary history or selective maintenance within the Atlantic subpopulation. Within the Atlantic subpopulation, we observed a distinct geographic and genetic structure in the distribution of B-positive genotypes (**Fig. 3B**). The B-Chromosome was identified predominantly in the Northern Coastal Plains (22.5%; 18 of 80 genotypes) and the Southern Coastal Plains (13.1%; 8 of 61), indicating that its presence is strongly associated with specific genetic lineages rather than being uniformly distributed across the range.

To reconstruct the evolutionary trajectory of these B-positive genotypes, we combined this population screen with a coalescent-aware phylogeny of the Atlantic subpopulation (**Fig. 3C**). The resulting topology places the Southern Coastal Plains carriers in a southern clade and the high-frequency Northern carriers in a distinct northern clade; although the CASTER tree is unrooted, the post-glacial northward expansion of the Atlantic subpopulation (Lovell et al. 2021) suggests that the southern clade represents the ancestral part of its range. The B-Chromosomes identified across this broad geographic and phylogenetic range, including those from the southern genotype J586A and the Chesapeake genotype J621C, share largely conserved gene content and nearly identical sequence where they align (∼1.4 SNPs per aligned kb; **Table S3**). This high degree of conservation is most simply explained by a single origin within the Atlantic lineage that predates the diversification of its present-day genetic groups. Its present-day distribution is therefore most consistent with persistence in the ancestral Atlantic subpopulation, represented by the southern clade, followed by northward dispersal via historical gene flow. The higher frequency of B-positive genotypes within the northern range (inset in **Fig. 3B**) could reflect demographic history and gene flow, context-dependent fitness effects, or both.

We next asked how often the progeny of B-positive plants are themselves B-positive. We used two independent genotyping markers with B-Chromosome-specific primers (**Table S13**) to screen the greenhouse-grown open-pollinated progeny of 23 B-positive maternal plants (all confirmed by *k*-mer screening; **Tables S14** and **S15**) using our primary primer (P001) and verified these calls with a secondary primer (P003). The two markers showed strong agreement (Cohen’s Kappa = 0.881, *p* < 0.001; 385 of 406 concordant calls, with all 21 discordant progeny P001-negative and P003-positive) (**Fig. S9**). A Generalized Linear Mixed Model (GLMM) with maternal family as a random effect estimated a B-positive frequency of 70.3% for a typical family (inverse-logit of the model intercept; 95% CI 56–81%; pooled proportion 266 of 406 = 65.5%) (**Fig. 3D**; **Table S15**). B-positive frequencies varied substantially among families (random-intercept variance = 1.91 on the logit scale; family proportions ranged from 0 to 100%, with 18 of 23 families above 50%). Because these families are open-pollinated, the pollen donors and their B-Chromosome status are unknown. Additionally, maternal plants may carry multiple B copies, as do both reference genotypes (about two in J621C and four in J586A; **Table S2**). While switchgrass is largely self-incompatible (Martínez-Reyna and Vogel 2002), any selfing or preferential mating between B-positive plants would also increase the proportion of B-positive offspring. This variation among families could reflect maternal B copy number, the pollen donors’ B status, or different B variants, similar to rye, where B accumulation is genetically controlled (Puertas et al. 1998). To separate these possibilities, future studies must use controlled reciprocal crosses with genotyped parents and paternity assignment.

### Transcriptional organization and functional profiling of B-linked genes

To investigate the transcriptional activity and regulatory coordination of B-linked genes, we performed Weighted Gene Co-expression Network Analysis (WGCNA) on the leaf tissue expression of B-linked genes across a set of carriers (B+) and non-carriers (B−) balanced in size and matched in genome-wide principal-component space (15 B+ and 15 B− genotypes, two replicate libraries each; **Table S16**). Of the 207 B-linked genes, 109 passed the expression filter, and WGCNA resolved them into a single coherent co-expression module (hereafter the B module; 92 genes) and 17 genes that were not assigned to any module because they lack consistent co-expression (**Fig. 4A**; **Table S17**). Genes in the B module showed coordinated expression that was consistently higher in B+ than in B− individuals, in both the Southern Coastal Plains (SP) and Northern Coastal Plains (NP) genetic subgroups (**Fig. 4B**). Normalized expression abundance [log₂ (TPM + 1)] further confirmed that genes of the B module are expressed at substantially higher levels in both SP B+ and NP B+ groups relative to non-carriers (**Fig. 4C**). The module eigengene correlated strongly with B-Chromosome presence (*r* = 0.80, *p* < 0.001) but negligibly with genetic group (*r* = 0.08) (**Fig. 4D**). Because the unassigned genes do not form a module (their first principal component explains 18% of their expression variance, versus 75% for the module), they were tested individually: none was associated with B status, and 5 differed in expression between the genetic groups (4 higher and 1 lower in NP; BH-adjusted *p* < 0.05) (**Fig. 4D**; **Table S17**). These results indicate that the expression of B-Chromosome genes is strongly associated with carrier status in the sampled individuals. GO enrichment of the B module (**Fig. 4E**) revealed significant enrichment (BH-adjusted *p* < 0.05) for microtubule-based movement, microtubule motor activity, microtubule binding, and the kinesin complex, alongside reciprocal meiotic recombination, the mismatch-repair complex, the mitotic cell cycle, the condensin complex, and chromosome segregation. Collectively, these findings suggest that the B-Chromosome maintains an organized transcriptional program enriched for functions that could support its own replication, repair, and segregation.

**Fig. 4.**
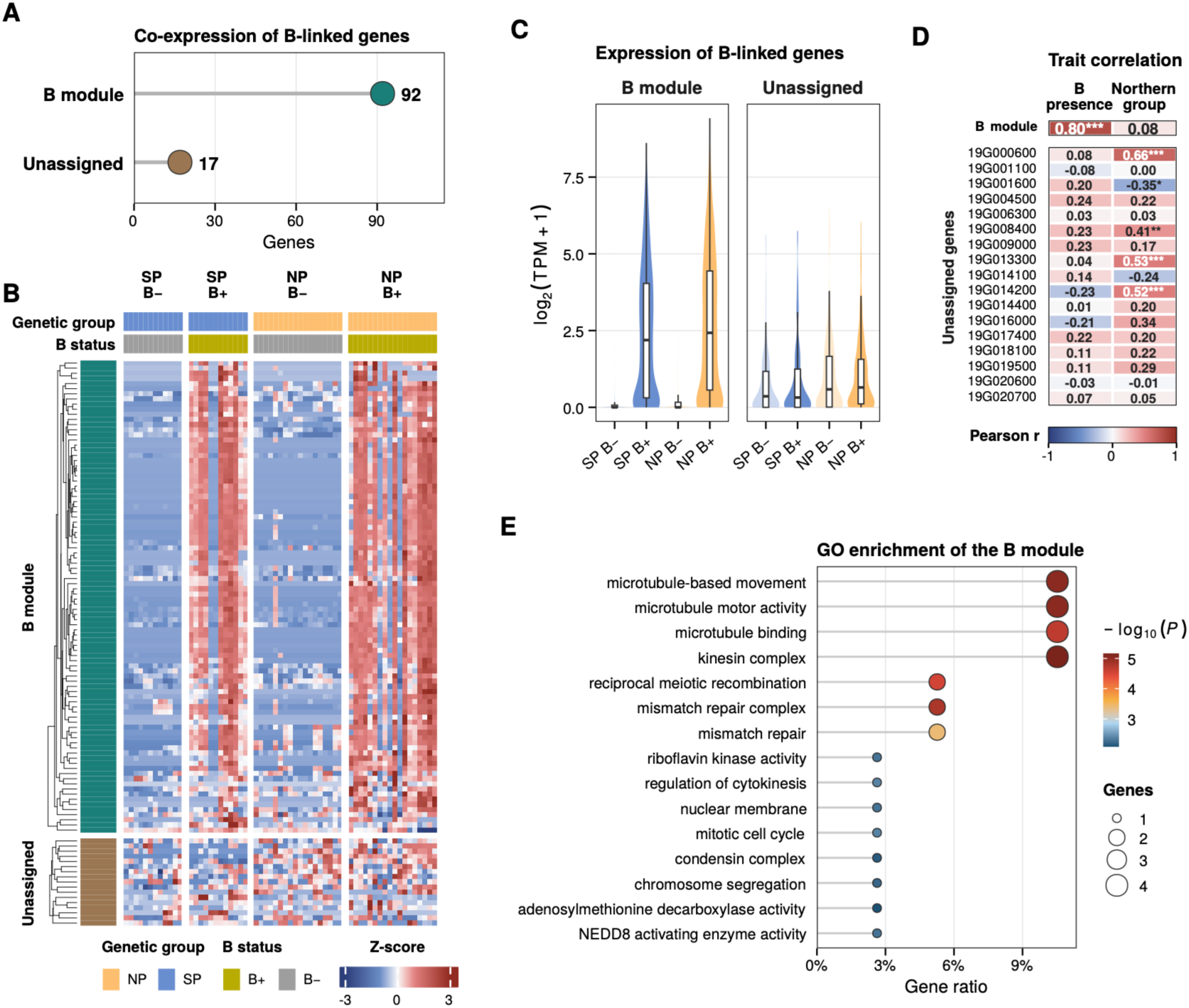
Transcriptional organization of B-Chromosome genes revealed by WGCNA. Analysis uses carrier (B+) and non-carrier (B-) individuals balanced in sample size. (**A**) Number of expressed B-linked genes in the single WGCNA co-expression module (B module, n = 92) and of genes not assigned to any module (unassigned, n = 17; brown). (**B**) Z-score expression heatmap of B-linked genes across all samples grouped by genetic group (SP, Southern Coastal Plains; NP, Northern Coastal Plains) and B-Chromosome status (B+, gold; B−, grey; colors as in **Figs. 1-3**). (**C**) Distribution of log₂(TPM + 1) expression (violin plots with box plots; boxes, median and interquartile range) of the genes of the B module and of the unassigned genes by genetic group and carrier status. (**D**) Top, Pearson correlation of the module eigengene with B-Chromosome presence and genetic group (NP); bottom, the unassigned genes, which do not form a module and are therefore not summarized by an eigengene, tested individually (Pearson r; BH-adjusted). Significance: \**p* < 0.05, \*\**p* < 0.01, \*\*\**p* < 0.001. (**E**) Gene Ontology (GO) enrichment of the B module. Dot size indicates gene count, and color indicates −log₁₀(*p*-value).

### B-presence-associated expression differences on the A-Chromosomes

To investigate whether B presence is associated with the expression of A-Chromosome genes, we performed a differential expression analysis comparing the A-Chromosome transcriptomes of B-positive (B+) and B-negative (B-) genotypes within the SP and NP genetic subgroups, respectively, using carrier and non-carrier sets balanced in size and matched in genome-wide principal-component space, with the leading principal components included as covariates (Methods) (SP: 6 B+ and 6 B− genotypes; NP: 9 B+ and 9 B− genotypes; two replicate libraries per genotype; **Table S16**). This comparison identified differentially expressed genes (DEGs) on the A-Chromosomes in both genetic subgroups (**Fig. 5A**; **Table S18**). In the SP background, we identified 94 upregulated and 47 downregulated DEGs in B+ individuals. In the NP background, we observed 121 upregulated and 18 downregulated DEGs. These findings suggest that B-Chromosome presence is associated with altered A-Chromosome gene expression. Additionally, we found that most of these transcriptional responses were subgroup specific. Among the upregulated DEGs, only one gene (<1% of the 214 upregulated genes) was shared between the SP and NP backgrounds, while 93 and 120 genes were exclusively upregulated in the SP and NP groups, respectively (**Fig. 5B**). Similarly, no downregulated DEG was shared, with 47 and 18 genes uniquely downregulated in the SP and NP groups, respectively (**Fig. 5C**). These results suggest that genetic background may influence how A-Chromosome expression differs with B-Chromosome presence, although the low overlap between two gene lists can also partly reflect limited statistical power or population structure confounding. The magnitude and significance of these changes are shown in **Fig. 5D-E**, and GO enrichment revealed distinct functional patterns in the two subgroups. In the SP background (**Fig. 5F**; **Table S19**), downregulated DEGs were enriched for heme transport, heme transmembrane transporter activity, the BLOC-1 complex, and glucose-1-phosphate adenylyltransferase activity. Upregulated SP DEGs showed no enriched GO term (smallest BH-adjusted *p* = 0.074). In the NP background, upregulated DEGs were primarily associated with photosynthesis (light reaction), terpene synthase activity, and lyase activity. In contrast, NP downregulated DEGs were strongly enriched for DNA metabolic processes, mismatch repair, mismatched DNA binding, and DNA topoisomerase type II activity, as well as glycosaminoglycan and heparan sulfate biosynthetic processes. These results indicate that B-presence-associated expression differences involve specific molecular pathways and differ between genetic backgrounds.

**Fig. 5.**
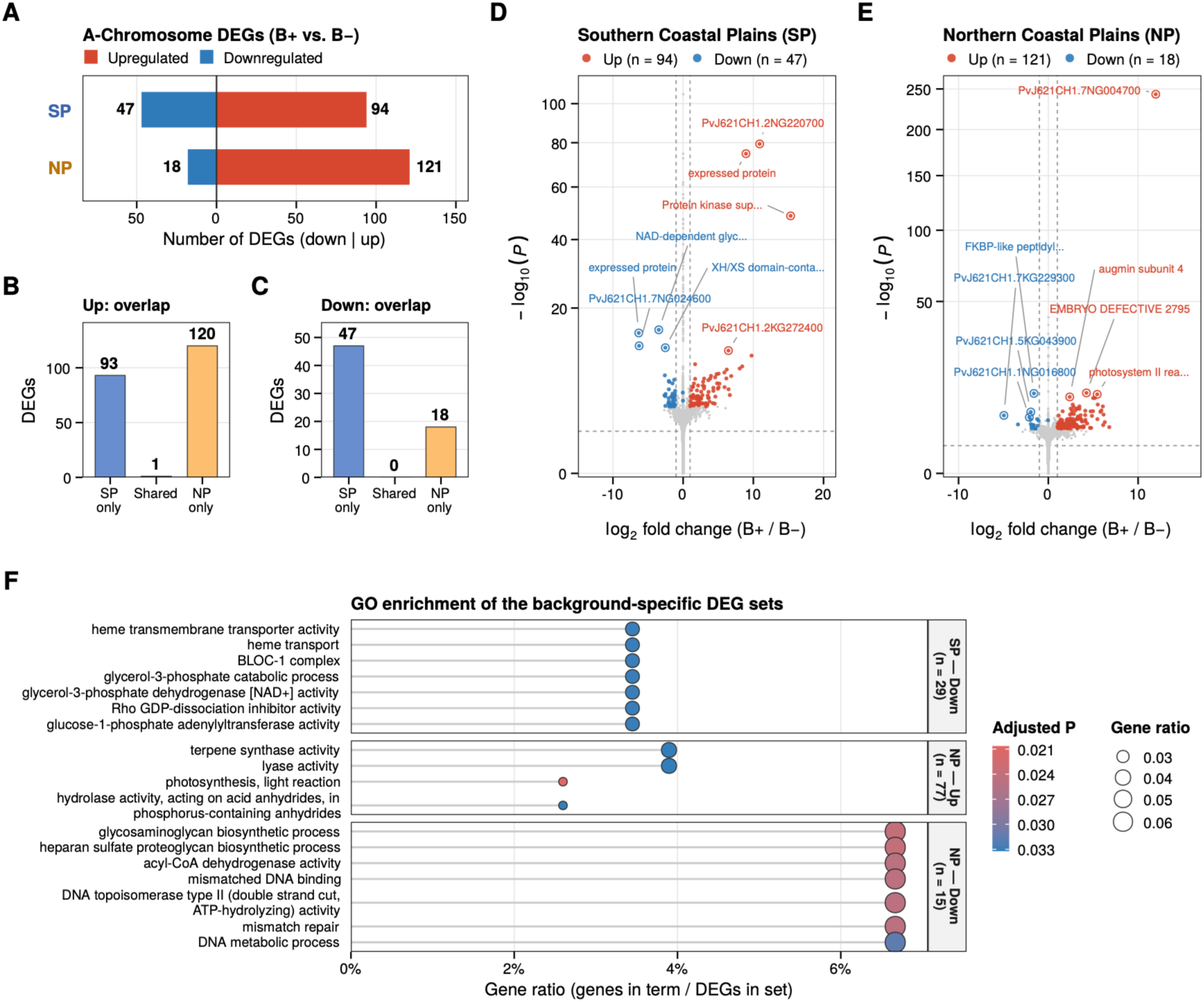
B-presence-associated A-Chromosome expression differences across two Atlantic groups. Differential expression compares B-positive (B+) to B-negative (B-) genotypes within the Southern Coastal Plains (SP) and Northern Coastal Plains (NP) groups using samples balanced and matched for genome-wide population structure (first three principal components as covariates). (**A**) Number of upregulated (right) and downregulated (left) differentially expressed genes (DEGs) located on the A-Chromosomes when comparing B-positive (B+) to B-negative (B-) genotypes within the SP and NP groups. (**B** and **C**) Number of upregulated (B) and downregulated (C) A-Chromosome DEGs found only in SP, in both groups (shared) or only in NP, showing limited overlap between the subgroup-specific DEG sets. (**D and E**) Volcano plots displaying the magnitude (log_2_ fold change) and significance (−log10 *p*-value) of A-Chromosome differential expression in the SP (**D**) and NP (**E**) groups. Dashed lines indicate the significance thresholds (*padj* < 0.05, |log_2_FoldChange| > 1). The four most significant upregulated and downregulated DEGs are labeled with their predicted function (gene identifier when no function is annotated). (**F**) GO enrichment analysis comparing the distinct molecular functions altered by B-Chromosome presence across the specific DEG sets (up to seven terms per set, those with the smallest adjusted p-values, ordered by gene ratio; number of DEGs with GO annotation in parentheses). Upregulated SP DEGs showed no enriched GO term (smallest BH-adjusted *p* = 0.074; **Table S19**). Dot size represents the gene ratio, and color gradient indicates the adjusted *p*-value.

## Discussion

B-Chromosomes have been recognized for more than a century and recorded in thousands of plant species, yet nearly all molecular insight into these elements derives from a few domesticated systems, principally maize and rye (Jones et al. 2008; Martis et al. 2012; Blavet et al. 2021). Our chromosome-level assemblies of two Atlantic switchgrass genotypes revealed a ∼34-Mb supernumerary B-Chromosome segregating in natural populations, to our knowledge the first supernumerary chromosome documented in *Panicum virgatum* and among the few complete B-Chromosome assemblies available for any wild plant. Switchgrass genomic studies anchored to the Gulf reference, from QTL mapping along a latitudinal gradient (Lowry et al. 2019) to range-wide population genomics (Lovell et al. 2021; Napier et al. 2022) had no capacity to detect this element; its independent recovery in genotypes from opposite ends of the Atlantic geographic range, corroborated by Hi-C contact maps and by long-read depth showing two to four copies per nucleus, establishes it as a genuine chromosome rather than an assembly artifact. The broader lesson concerns reference bias. When a single genotype stands in for a species, presence-absence variation private to diverged lineages, here, an entire chromosome, remains invisible, leaving analyses of expression, structure, and adaptation conditioned on an incomplete genome. The similar recovery of an unsuspected B-Chromosome from a telomere-to-telomere assembly of the wild grass *Phragmites australis* (Cui et al. 2025) suggests that lineage-resolved and pangenome-scale sequencing will continue to surface accessory chromosomes that cytology-based catalogs undercount (D’Ambrosio et al. 2017).

The architecture of the Atlantic B-Chromosome identifies it as a product of the A-Chromosome complement, assembled by rearrangement rather than inherited intact. Diagnostic *k*-mers and gene synteny trace the element to fragmented blocks of at least four autosomes (Chr01, Chr02, Chr07, and Chr09) drawn from both subgenomes, with most of the subgenome-assigned sequence attributed to the N subgenome. The element is also markedly bipartite: a gene-rich, largely single-copy proximal region (∼8 Mb) adjoins a repeat-dominated, gene-poor distal region (∼27 Mb) rich in LTR/Ty3 elements and punctuated by Mutator TIR peaks. This anatomy closely parallels the two best-characterized plant B-Chromosomes: the rye B is a mosaic of A-Chromosome fragments and organellar DNA (Martis et al. 2012), and the maize B couples a small genic fraction to massive repeat accumulation (Blavet et al. 2021; Liu et al. 2025); and the 207 protein-coding genes retained on the switchgrass B are similar in number to the 229 and 277 genes reported for the *Aegilops speltoides* and *P. australis* Bs (Ruban et al. 2020; Cui et al. 2025). Whether the switchgrass B, like the rye B, also carries organelle-derived sequence has not yet been tested. This convergent structure is consistent with models in which B-Chromosomes originate through chromoanagenesis-like shattering and reassembly of A-Chromosome fragments, plausibly triggered by hybridization or polyploidization, and then degenerate as repeats accumulate on a chromosome that rarely recombines (Johnson Pokorná and Reifová 2021; Houben et al. 2025). Because the element carries material from both subgenomes, it most plausibly formed after their union in the allotetraploid ancestor (≤4.6 MYA; Lovell et al. 2021). The strong N-subgenome bias remains unexplained. It could reflect greater instability of N-subgenome chromosomes when the B formed, homology-mediated capture of N-subgenome sequence during later rearrangements, or preferential retention of N-derived segments. These alternatives differ in timing: segments present since the origin of the B should be as old as its oldest sequence, whereas segments captured later should be younger, so dating the N- and K-derived segments (for example, from the insertion ages of the LTR retrotransposons they contain) and examining the junctions between donor segments could distinguish them.

The persistence of the element is as remarkable as its origin. Intact LTR retrotransposons on the B are younger than those on any A-Chromosome (median 335 versus 473-720 kyr). Their ages tend to be higher in the gene-rich proximal region (median 667 kyr) than in the repeat-rich distal region (283 kyr), which would fit a proximal region that retains sequence inherited from its A-Chromosome donors and a distal region that grew by recent accumulation of LTR/Ty3 elements, although the number of intact elements in each region is modest. A deficit of old intact elements could also arise if older LTRs decay or are removed faster on the B, so these ages are best read as describing the turnover of its repeat complement. These insertion ages date repeat accumulation rather than the formation of the B and depend on the assumed substitution rate; nevertheless, a repeat complement younger than that of any A-Chromosome fits a recent history of the B, consistent with its restriction to the Atlantic lineage. B-Chromosomes with conserved gene content and shared diagnostic sequences occur in both the southern and the northern clades, despite structural polymorphism among the assembled copies. This distribution is most parsimoniously explained by a single origin within the Atlantic lineage, after its divergence from the Gulf and Midwest subpopulations (∼0.7-1.4 MYA; Lovell et al. 2021), followed by persistence and redistribution through gene flow. Persistence of a largely repetitive, gene-poor element poses the classical B-Chromosome paradox: such an element should be progressively eliminated unless it benefits the plants that carry it or is transmitted at higher-than-Mendelian rates (Östergren 1945; Jones 1995; Camacho et al. 2000; Ågren and Clark 2018), and both forces can operate at once, as shown for centromere-associated meiotic drive opposed by female fitness costs in *Mimulus* (Fishman and Kelly 2015). Our data do not yet distinguish among these possibilities. The Atlantic B occurs in multiple copies (about two per nucleus in J621C and four in J586A), a variation in number between individuals that is typical of B-Chromosomes, reaches appreciable, geographically structured frequencies (22.5% and 13.1% of genotypes in the Northern and Southern Coastal Plains) while being rare among Gulf and absent among Midwest genotypes surveyed, and is present in most open-pollinated progeny of B-positive plants (70.3% for a typical family). This progeny frequency, however, is also expected from multiple maternal B copies, B-positive pollen donors, or occasional selfing, so it does not by itself demonstrate biased transmission.

In cereals, B-Chromosomes are known to accumulate through directed nondisjunction at gametophytic mitoses, the first pollen mitosis in rye, the second in maize, coupled there with preferential fertilization, under the control of B-encoded factors (Banaei-Moghaddam et al. 2012; Birchler and Yang 2021; Chen et al. 2022); comparable nondisjunction, accompanied by an extra pollen mitosis, was recently documented for the B-Chromosome of a wild sorghum (Karafiátová et al. 2024). In rye the drive determinant has now been identified, *DCR28*, a serially amplified, B-encoded microtubule-associated protein expressed at first pollen mitosis (Chen et al. 2024), and a kinesin motor likewise underlies the independent meiotic drive of maize abnormal chromosome 10 (Dawe et al. 2018). Against this backdrop, the transcriptional organization of the switchgrass B is compatible with, but does not demonstrate, a drive mechanism. Its expressed genes form a single coherent module whose activity tracks B presence (*r* = 0.80) but not host genetic subgroup, and that is enriched for microtubule motor, kinesin, condensin, cell-cycle, and chromosome-segregation functions, the functional classes from which known drive factors have repeatedly emerged. The B itself carries four kinesin motor-domain genes, all in this module and expressed predominantly in B-positive plants (**Table S4**). None of these observations distinguishes drive from selection: a B that benefits its carriers would show the same population frequencies and the same carrier-specific expression. Whether the switchgrass B accumulates by such a mechanism is untested: our progeny data are frequencies of B-positive offspring in open-pollinated families, which cannot be converted into transmission rates without known pollen donors, parental B copy numbers, and controlled reciprocal crosses. The substantial among-family variation in B-positive frequency (variance 1.91 on the logit scale) could reflect maternal B copy number, the B status of pollen donors, maternal genetic background, or structural variants among B copies, as well as heritable variation of the kind documented for B accumulation in rye (Puertas et al. 1998); controlled crosses combined with cytological observation of gametophytic divisions could resolve these alternatives.

The B-Chromosome may also influence the expression of A-Chromosome genes: B presence was associated with differential expression of 141 A-Chromosome genes in the Southern background (94 upregulated, 47 downregulated) and 139 in the Northern background (121 upregulated, 18 downregulated), an excess of upregulation in both (67% and 87% of DEGs, respectively). Several mechanisms, which are not mutually exclusive, could produce such differences: genes carried on the B could regulate A-Chromosome genes directly; reads from B-linked genes could be mis-assigned to their A-Chromosome paralogs; and the extra chromatin could alter the balance of dosage-sensitive regulatory complexes, a general consequence of changing chromosome dosage (Birchler and Veitia 2012; Shi et al. 2021); dosage- and tissue-dependent effects of B-Chromosomes on A-Chromosome expression are likewise documented across the maize life cycle (Huang et al. 2016; Hloušková et al. 2025; Liu et al. 2025). These mechanisms describe how expression differences could arise, not whether they affect fitness, and any of them could be selectively neutral. These associations are, however, fundamentally observational. B status remains partially entangled with fine-scale population structure, and the sequence homology between B-linked genes and their A-Chromosome progenitors can generate cross-mapping artifacts. While our balanced sampling and covariate modeling mitigate these analytical hurdles, isolating the true regulatory footprint of the B-Chromosome will ultimately require comparing segregating siblings within controlled crosses using strictly uniquely mapping reads.

The clearest feature of these expression differences is their dependence on genetic background. Of the 279 distinct A-Chromosome genes called differentially expressed in either background (141 in the Southern and 139 in the Northern background), only one (<1%) was shared between them, and the enriched functions diverged accordingly: in the Southern background only the downregulated set was enriched, for heme transport, the BLOC-1 complex and carbohydrate-metabolic activities, whereas in the Northern background upregulated genes were enriched for photosynthesis and terpene synthase activity and downregulated genes for DNA metabolism, mismatch repair and topoisomerase II activity. The same accessory chromosome is thus associated with different expression changes in different genetic backgrounds, although low overlap between two gene lists can also arise when statistical power is limited, so formal tests of a B-by-background interaction in larger samples are needed. Background- and dosage-dependent effects have long been suspected from phenotypic studies of plant B-Chromosomes (Bougourd and Jones 1997) and are increasingly visible transcriptomically (Hloušková et al. 2025). Whether such context dependence extends to fitness, and hence shifts the balance between the processes that raise and lower B frequency (Camacho et al. 2000; Johnson Pokorná and Reifová 2021), remains to be tested. The uneven geographic distribution we observe could accordingly reflect demographic history and gene flow, background-dependent fitness effects, or both; our data do not yet discriminate among these.

Whether the switchgrass B is costly, neutral, or conditionally beneficial to the plants that carry it is the central open question; we have not yet measured its fitness effects, and it may be maintained by a balance of opposing forces rather than by any single one. Natural populations are the appropriate arena in which to answer it. Classic studies of *Allium schoenoprasum* demonstrated that B frequencies in the wild can track selection on carriers, including superior germination of B-carrying seed under drought (Holmes and Bougourd 1989; Plowman and Bougourd 1994). The resources established here provide a foundation for testing how the fitness of B-positive plants and the frequency of B-positive progeny shape its persistence in natural populations. More broadly, our results add whole accessory chromosomes to the inventory of standing structural variation segregating in natural plant populations; a cryptic axis of genomic diversity, invisible to short-read and single-reference approaches, with potential consequences for gene dosage, recombination, and genome evolution (Johnson Pokorná and Reifová 2021; Ferree et al. 2024). With a complete B sequence, population-scale B-status data, progeny families, and established common gardens, Atlantic switchgrass now provides a tractable system for moving the study of plant B-Chromosomes from domesticated fields into the ecological and evolutionary settings in which these elements evolve.

## Materials and Methods

### Plant material

For reference genome assembly, two genotypes from the Atlantic subpopulation, J621C and J586A, were selected for de novo genome sequencing. J621C (northern/Chesapeake-associated) was the primary Atlantic reference; J586A (southern) provided an independent, geographically distinct assembly and a second B carrier used to select conserved regions for PCR primer design. For the population-genomic analysis, we used Atlantic genotypes previously described in (Lovell et al. 2021), representing a broad geographic and genetic sampling of the Atlantic coast. Because the analyses focus on natural populations, genotypes originating from breeding populations (breeding selections, n = 11) or cultivars (n = 12) were removed using the collection-type records, leaving 280 natural-collection genotypes (**Table S9**).

To estimate the frequency of B-positive progeny, 23 open-pollinated families were collected from common-garden nurseries, with maternal parents selected from B-positive Atlantic individuals; 406 F1 progeny from these 23 families (8-20 progeny per maternal line; mean 17.7) were grown under greenhouse conditions for PCR screening (**Tables S14** and **S15**).

### Library preparation and sequencing

We sequenced the two genotypes of the Atlantic subpopulation (J621C and J586A) using a whole genome shotgun sequencing strategy and standard sequencing protocols. Sequencing reads were collected using Illumina and PacBio platforms. Illumina and PacBio reads were sequenced at the HudsonAlpha Institute in Huntsville, Alabama. Illumina reads were sequenced using the Illumina NovaSeq 6000 platform, and PacBio HiFi (circular consensus) reads were generated on the PacBio REVIO platform (four SMRT Cells per genotype; 117.2 Gb of HiFi reads for J621C and 103.3 Gb for J586A; **Table S2**). For each genotype, one 400bp insert 2×150 Illumina fragment library was sequenced along with one 2×150 Omni-C library. Prior to assembly, Illumina fragment reads were screened for PhiX contamination. Reads composed of >95% simple sequence were removed. Illumina reads <50bp after trimming for adapter and quality (q<20) were removed. The coverage of each library type (Illumina, Omni-C, PacBio) for each genotype is detailed in **Table S1**.

### Genome assembly and construction of pseudomolecule chromosomes

Each genome assembly was generated by assembling the PacBio HiFi reads using hifiasm in Hi-C mode (Cheng et al. 2021) and subsequently polished using RACON (Vaser et al. 2017). This produced initial haplotype assemblies denoted as HAP1 and HAP2. Assembly sizes, contig N50, etc. can be found in **Table S1** for each genotype.

Omni-C Illumina reads from each *Panicum virgatum* genotype were aligned to their respective initial contig set with bwa mem (Li 2013), and chromosome scale scaffolding was performed using a combination of the Hi-C read alignments and the Juicebox contact map viewer (Robinson et al. 2018). Each assembly haplotype bin was inspected for misjoins and used to make additional joins among contigs to generate the chromosome assemblies described below: 19 chromosomes in J621C HAP1, J621C HAP2, and J586A HAP1, and 18 in J586A HAP2. Chromosomes were ordered and oriented using version 5.0 release of *Panicum virgatum* var. AP13. Each chromosome join is padded with 10,000 Ns. Contigs terminating in significant telomeric sequence were identified using the (TTTAGGG)n repeat, and care was taken to make sure that they were properly oriented in the production assembly. The remaining scaffolds were screened against bacterial proteins, organelle sequences, GenBank nr and removed if found to be a contaminant. After forming the chromosomes, it was observed that some small (<20Kb) redundant sequences were present on adjacent contig ends within chromosomes. To resolve this issue, adjacent contig ends were aligned to one another using BLAT (Kent 2002), and duplicate sequences were collapsed to close the gap between them. Total misjoins, joins and contig collapses for each haplotype assembly can be found in **Table S1**.

Last, homozygous SNPs and INDELs were corrected in each respective genome by aligning Illumina reads (2×150, 400bp insert) using bwa mem (Li 2013) and identifying homozygous SNPs and INDELs with the GATK’s UnifiedGenotyper tool (McKenna et al. 2010). Homozygous SNPs and INDELs were corrected in each haplotype assembly. Final genome statistics for each assembly (Homozygous SNPs and INDEL fixes, contig N50, genome assembly sizes, etc.) can be found in **Table S1**.

### Genome quality assessment

Completeness of the euchromatic portion of each assembly was assessed by aligning the version 5.0 *P. virgatum* var. AP13 primary annotated transcripts onto each finalized genome assembly using blat. Genes that aligned at greater than 90% identity and 85% coverage were retained. The aim of the completeness analysis is to obtain a measure of completeness of the assembly, rather than a comprehensive examination of gene space. Final alignment statistics for each assembly can be found in **Table S1**.

### Genome annotation

A repeat library was created from de novo repeats predicted by RepeatModeler2 (Flynn et al. 2020) on the *Panicum virgatum* genome. The predicted repeats underwent functional analysis through InterProScan (Jones et al. 2014), incorporating the Pfam (Mistry et al. 2021) and PANTHER (Mi et al. 2019) databases. Any repeats that displayed significant hits to protein-coding domains were subsequently excluded from the repeat library. Finally, the constructed species-specific repeat library was used to soft-mask the *Panicum virgatum* genomes with RepeatMasker (Smit et al. 2013–2015).

Transcript assemblies used for annotation were made from 150 bp stranded paired-end Illumina RNA-seq reads pooled across a diverse panel of tissues and developmental stages, using PERTRAN, which conducts genome-guided transcriptome short read assembly via GSNAP(Wu and Watanabe 2005) and builds splice alignment graphs after alignment validation, realignment, and correction. Putative full-length transcripts were generated from PacBio Iso-Seq. Circular Consensus Sequences (CCSs) were corrected and collapsed by a genome-guided correction pipeline. The pipeline aligns CCS reads to the genome with GMAP (Wu and Nacu 2010), corrects small indels in splice junctions, and clusters alignments when all introns are the same or >= 95% overlap for single-exon alignments. Subsequently, PASA (Haas et al. 2003) was used to construct transcript assemblies by combining the transcript assembly sets described above.

Putative gene loci were determined by transcript assembly alignments and/or EXONERATE (Slater and Birney 2005) alignments of proteins from *Paspalum vaginatum*, *Panicum hallii*, *Zea mays*, *Sorghum bicolor*, *Setaria viridis*, *Oryza sativa*, *Pharus latifolius*, *Chasmanthium laxum*, *Brachypodium stacei*, *Joinvillea ascendens*, *Acorus americanus*, *Asparagus officinalis*, *Ananas comosus*, *Musa acuminata*, *Elaeis guineensis*, *Populus trichocarpa*, *Aquilegia coerulea*, *Arabidopsis thaliana*, *Glycine max*, *Vitis vinifera*, and Swiss-Prot eukaryote proteomes to repeat softmasked *Panicum virgatum* genomes with up to 2K BP extension on both ends unless extending into another locus on the same strand. Gene models in each locus were predicted by homology-based predictors, FGENESH+ (Salamov and Solovyev 2000), FGENESH_EST, EXONERATE, PASA assembly ORFs, and AUGUSTUS (Stanke et al. 2006) trained on the high confidence PASA assembly ORFs and with intron hints from short read alignments. The best-scored predictions for each locus were selected using multiple positive factors including EST and protein support, and one negative factor: overlap with repeats. The selected gene predictions were improved by PASA. The improvement included adding UTRs, splicing correction, and adding alternative transcripts.

PASA-improved gene model proteins were subject to protein homology analysis to the above-mentioned proteomes to obtain a Cscore and protein coverage. Cscore is a protein BLASTP score ratio to the mutual best hit BLASTP score and protein coverage is the percentage of protein aligned to the best of homologs. PASA-improved transcripts were selected based on Cscore, protein coverage, EST coverage, and their CDS overlap with repeats. The transcripts were selected if their Cscore and protein coverage were >= 0.5 or if covered by ESTs. For gene models whose CDS were overlapped by repeats by more than 20%, their Cscore had to be at least 0.9 and homology coverage at least 70% to be selected. The selected gene models were subject to Pfam analysis and gene models without strong transcriptome and homology support whose proteins were more than 30% overlapped by Pfam TE domains were removed. Incomplete gene models, low homology supported without fully transcriptome-supported gene models, short single exon (< 300 BP CDS) without protein domains nor good expression, and repetitive gene models without strong homology support were manually filtered out.

### Comparative synteny analyses

Syntenic relationships among five assemblies (AP13 v5.1 and the two haplotypes of J586A and J621C) were inferred with GENESPACE (Lovell et al. 2022), which combines OrthoFinder orthogroups (Emms and Kelly 2019) with collinearity-constrained syntenic block detection, and were plotted as a riparian plot (**Fig. 1**). Gene-level synteny between the B-Chromosome and the A-Chromosomes of J621C, AP13, VS16 and *Panicum hallii* (**Fig. S6**) was obtained with the MCscan pipeline in JCVI (Tang et al. 2008): primary-transcript CDS were compared with *jcvi.compara.catalog* ortholog (C-score ≥ 0.99), anchors were filtered with *jcvi.compara.synteny* screen (minimum span of 30 genes for the genome-wide comparisons) and visualized with *jcvi.graphics.karyotype*.

### B-Chromosome repeats landscape, centromere identification, sequence origin, and age estimation

EDTA (Ou et al. 2019) was used to annotate the J621C repeat landscape. Intact-repeat GFF3 annotations were combined with primary-transcript (gene) coordinates from the gene annotation, and each base of B1 and of the A-Chromosomes was assigned to at most one class (genes first, then LTR/Ty3, LTR/Ty1, Helitron, Mutator, CACTA, hAT, PIF/Harbinger, Tc1/Mariner and LTR/unknown elements), so that overlapping annotations are counted once. The annotated fraction of each class was summarized in 500-kb windows moved in 50-kb steps (**Fig. 2A** for Chr19; **Fig. S5** for the 18 A-Chromosomes) and over each whole chromosome as a share of its annotated sequence (**Fig. 2B**). LTR/Ty1 (Copia), hAT, PIF/Harbinger and Tc1/Mariner elements, which make up similar small shares of the A- and B-Chromosomes, are shown together as other TEs. TRASH (Wlodzimierz et al. 2023) identified a candidate 166-bp tandem-repeat motif abundant near the middle of A-Chromosomes, particularly in the K subgenome. The motif was decomposed into 24-mers and used to mask J621C HAP1 Chr19. Masked bases were summarized in non-overlapping 50-bp windows. Runs were initiated by at least 10 windows with more than 90% masking, with a reported tolerance of up to five intervening windows. This procedure yielded the two candidate intervals reported in Results. For sequence assignments, the 8,181,153 discoverable B1 single-copy markers were treated as single-end reads and aligned with bwa-mem2 (Vasimuddin et al. 2019). The top A-Chromosome alignment for each marker was projected onto its B1 coordinate. Markers were ordered by their B1 coordinate, consecutive markers with the same best-hit autosome (or subgenome) were grouped into runs, and runs of at least 200 markers were retained; the sequence assigned to each autosome or subgenome is the summed span of its runs (**Table S7**). Intact LTR insertion ages were estimated from paired-LTR divergence as T = d/2r, where d is the divergence between the two LTRs and r = 1.5 × 10⁻⁸ substitutions per site per year (Koch et al. 2000). Median ages were summarized chromosome-wide (with bootstrap standard errors, 2,000 resamples), for the proximal (<8 Mb) and distal (≥8 Mb) regions of the B, and in 500 kb windows with a 10 kb step. Ages assume a uniform substitution rate and are not corrected for gene conversion between LTRs, so they are relative estimates. Repeat composition, candidate centromeric intervals, *k*-mer assignments and LTR ages are provided in **Tables S5-S8**.

### Population-level *k*-mer screening for B-Chromosome presence

To identify individuals in the resequencing panel that carry the B-Chromosome we defined a set of single-copy 80-mers unique to the B. First, the assemblies of AP13, DAC6, VS16, WBC, J386C, J514A, J610B and J621C (A-Chromosomes only, n = 18 per genotype) were decomposed into 80-mers. Second, Chr19 of J621C (HAP1) was decomposed into 80-mers, retaining only those present in a single copy on the B. Third, the combined autosomal 80-mer set was subtracted from the single-copy B set, leaving B-Chromosome markers absent from the screened A-Chromosome assemblies. Markers were counted in each resequenced genotype’s FASTQ library and scored as present if detected in at least one Illumina read. The J621C positive-control library defined 8,181,153 discoverable markers. Subpopulation frequencies were estimated in the 732-genotype tetraploid diversity panel of Lovell et al. (2021), of which 696 genotypes had a screened library, grouping genotypes by their subpopulation assignment in that study (Gulf, including Texas; Midwest; Atlantic) (**Tables S11** and **S12**). Each library was scored by the percentage of these markers present in its reads (**Table S11**; present = 100% − missing). The distribution was bimodal: 79.6% of libraries contained <5% of the markers, no library contained between 22.8% and 66.8%, and B-positive libraries (>60% present) contained 67–100%. Libraries with ≤10% of the markers present were classified as B-negative; libraries with intermediate values (10–23% present, mostly lower-coverage libraries; 8 of the 280 Atlantic genotypes) were excluded from frequency estimates (**Table S12**).

### SNP calling for the Atlantic subpopulation using J621C as a reference

A total of 324 genotypes from the Atlantic subpopulation diversity panel were sequenced using Illumina HiSeq X10 and Illumina NovaSeq 6000 paired-end sequencing at HudsonAlpha Institute for Biotechnology and the Joint Genome Institute. Raw sequencing reads are available under the NCBI SRA identifiers associated with Lovell *et al*. (2021; Supplemental Data 4). Those genotypes were selected based on the demographic history and previous population genetic structure analysis (Lovell et al. 2021). Breeding selections (n = 11) and cultivars (n = 12) were excluded to restrict the analyses to natural populations, and 280 natural-collection genotypes were retained for SNP calling and all population-genomic analyses (**Table S9**). Raw sequencing reads were mapped to the Atlantic J621C reference genome using BWA-MEM with default parameters in BWA v0.7.15 (Li and Durbin 2010). Aligned reads were filtered for duplicates and realigned using GATK v4.3.0 (DePristo et al. 2011). The filtered reads were sorted using SAMtools v1.8 (Li et al. 2009). PCR duplicates were marked using MarkDuplicate and AddOrReplaceReadGroup implemented in GATK v4.3.0 (DePristo et al. 2011) and filtered out using Picard (https://github.com/broadinstitute/picard). Genotypes of each individual were called using MPILEUP implemented in BCFTOOLS (minimum coverage of eight and a minimum alternate allele count of four) and then merged into a VCF file. Subsequently, the 24mer softmasked repeats with 25bp buffer around the repeats were removed, and only biallelic SNPs with <= 20% missing data and minor allele frequencies > 0.05 were retained. The remaining set of SNPs was then slightly linkage-disequilibrium-pruned to remove redundant SNPs using a threshold of r <= 0.95 across the whole genome in PLINK2.0 (Chang et al. 2015).

### Discovery, validation and comparison of the B-Chromosome

A single B-Chromosome was noted in the combined HAP1/HAP2 assembly of each genotype: for J621C, a complete 34.8-Mb copy on HAP1 and a shorter 15.7-Mb copy on HAP2; for J586A, a single 29.6-Mb copy on HAP1. The B is included as Chr19 in the HAP1 releases of both genotypes and in the J621C HAP2 release; the J586A HAP2 release contains the 18 autosomes only. Hi-C alignments and Juicebox contact maps were re-examined for each genotype and showed that the B forms a discrete contact domain with no detectable contact to any of the 18 autosomes (both subgenomes), excluding a technical or assembly artifact (**Fig. S1**). B-Chromosome copy number was estimated from HiFi read depth (see below; **Fig. S1C**; **Table S2**). The three assembled B copies (B1, B2, B3) were compared by whole-sequence alignment with minimap2 (Li 2018) followed by structural-variant annotation with SyRI (asm5 preset for B1–B2 and asm20 for the cross-genotype pairs) and visualization with plotsr (Goel et al. 2019; Goel and Schneeberger 2022) (**Fig. S2**; **Table S3**), and independently by FlexiDot (Seibt et al. 2018) dot plots (**Figs S3** and **S4**); and gene content was compared by reciprocal best BLASTP hits and OrthoFinder (**Table S3**). All downstream analyses use the complete B1 (J621C HAP1).

### B-Chromosome copy number from HiFi read depth

For each genotype, PacBio HiFi reads (four SMRT Cells; **Table S2**) were aligned with minimap2 (Li 2018) to a combined reference containing both haplotype assemblies and all assembled B copies (J621C: HAP1 + HAP2, including B1 and B2; J586A: HAP1 + HAP2, including B3), with each B copy included once. Because the reference contains both haplotypes, reads are placed on the haplotype from which they derive, and each A-Chromosome sequence represents one copy per diploid (2n) nucleus. Alignments were sorted, merged and indexed with SAMtools (Danecek et al. 2021), and read statistics were obtained with SeqKit (Shen et al. 2016). Read depth was computed in non-overlapping 100-kb windows with tinycov v0.4.0 (https://github.com/cmdoret/tinycov) and independently with mosdepth (Pedersen and Quinlan 2018) using all alignments and only alignments with mapping quality ≥20; window depths from the two tools were virtually identical (Pearson *r* > 0.9999). Windows with >10% assembly gaps and sequences shorter than 1 Mb were excluded. The single-copy depth (D_A) was defined as the median depth of all A-Chromosome windows of both haplotypes, and B copy number per 2n nucleus as the median depth of the B windows divided by D_A; 95% confidence intervals were obtained by block bootstrapping (2,000 replicates; blocks of ten consecutive windows) of the A and B windows. As a robustness check, copy number was recomputed from windows in which ≥90% of the depth came from uniquely placed reads (mapping quality ≥20). For J621C, B2 was aligned to B1 with minimap2, and B1 windows were classified as B1-specific (≤10% covered by B2 alignments) or shared (≥90% covered); because reads from shared sequence are distributed between the two near-identical copies, copy number in the shared region was estimated from the summed depth on B1 and B2. The median per-read divergence from the assembly was 0.10% for J621C and 0.13% for J586A, consistent with the reads deriving from the assembled plants (**Table S2**).

### DNA extraction from F1 progeny

High-quality genomic DNA was extracted from young leaves of switchgrass F1 progeny using a modified CTAB protocol. Leaf tissues were ground sufficiently, then combined with 800 μL of 2% CTAB buffer and 4 μL of RNase A, mixed thoroughly, and incubated at 65 °C for 30 min. Following centrifugation at 12,000 rpm for 10 min, the supernatant was collected. Subsequently, an equal volume of chloroform/isoamyl alcohol (24:1) was added, followed by centrifugation at 12,000 rpm for 10 min and retention of the supernatant. Pre-chilled isopropanol (4 °C) was then added in equal volume, mixed by inversion, and allowed to precipitate at -20°C for 60 min. The DNA precipitate was pelleted by centrifugation at 12,000 rpm for 15 min, washed twice with 500 μL of 70% ethanol, and dried at 37°C. Finally, the DNA was resuspended in water, incubated at 37 °C for 10 min, and stored at -20°C. DNA quality and quantity were assessed using a NanoDrop spectrophotometer.

### PCR-based screening of F1 progeny for B-Chromosome presence

To develop a PCR-based assay for detecting the B-Chromosome, we first identified DNA sequences that were both unique to the B-Chromosome and conserved across different Atlantic switchgrass individuals harboring it. For this, we utilized a comparative *k*-mer analysis between the B-Chromosome assembly (Chr19) of our primary reference genotype, J621C, and the whole genome assembly of a second B-Chromosome-carrying Atlantic genotype, J586A. Specifically, we extracted all unique (single-copy) 80-mers from the J621C Chr19 sequence and mapped these against the complete J586A genome assembly. We considered *k*-mers B-Chromosome-conserved if they mapped uniquely to Chr19 in J586A, showing minimal or no significant matches to the A-Chromosomes (Chr01-Chr18) in J586A. Based on this mapping, we identified contiguous regions of B-Chromosome conservation by merging the coordinates of qualifying 80-mers on the J621C Chr19 reference. This stringent merging process yielded a set of high-confidence B-Chromosome-specific sequence regions shared between the two genotypes. From this set, we selected regions ≥ 1000 bp in length with a GC content between 30% and 70% as candidates for primer design. If more than ten such regions met these criteria, we randomly chose ten for further processing.

Next, we designed primer pairs flanking these selected B-Chromosome-specific regions using the Primer3-py library (v2.6.1) (Koressaar and Remm 2007; Untergasser et al. 2012). We applied standard parameters targeting an amplicon size of 200–300 bp, a primer length of 22–27 nt (optimum 24 nt), a melting temperature (Tm) between 55-62°C, a maximum mononucleotide run of 4 and a GC clamp of 3, returning up to 5 primer pairs per region. To assess the specificity of the designed primers in silico, we individually searched each forward and reverse primer sequence against indexed BLAST databases created from the J621C B-Chromosome (Chr19) FASTA file and a separate FASTA file containing only the J621C A-Chromosomes (Chr01-Chr18). We conducted these searches using blastn with parameters optimized for short sequences (-task blastn-short, -evalue 0.1, -word_size 7) and parsed the output to identify potential binding sites. We selected primer pairs for experimental validation based on the following in silico criteria: (1) prediction of a single PCR product within the target size range (200-300 bp) on the B-Chromosome sequence, inferred from appropriately oriented forward and reverse primer hits; (2) ideally, only one high-identity hit for each primer within its target B-Chromosome region; and (3) prediction of zero amplicons on the A-Chromosomes. The final selected primer sequences are listed in **Table S13**.

Using the validated primer pairs, we screened the genomic DNA extracted from the 406 F1 progeny and parental controls via PCR. Thermocycling conditions were as follows: initial denaturation at 95°C for 15 min; followed by 35 cycles of denaturation at 95°C for 1 min, annealing adjusted based on primer Tm for 30 s, and extension at 72°C for 30 s; and a final extension at 72°C for 10 min. PCR products were visualized by electrophoresis on a 1% agarose gel stained with SYBR Safe run at 120 V for 45 min. We scored each F1 individual for the presence or absence of the B-Chromosome-specific amplicon based on the visualization of a band of the expected size, allowing us to score each progeny as B-positive or B-negative.

To ensure the accuracy of our presence/absence scoring, the F1 progeny were independently genotyped using both a primary primer (P001) and a secondary validation primer (P003). The reliability and concordance between these two independent markers were quantified using Cohen’s Kappa statistic. To estimate the average frequency of B-positive progeny and its variation among families, we employed a Generalized Linear Mixed Model (GLMM) implemented in R. The B-Chromosome status of each F1 individual was modeled as a binary response variable (scored B-positive only when both markers amplified; binomial distribution with a logit link function). To account for differences among maternal plants and to quantify among-family variability, the 23 distinct maternal families were incorporated into the model as a random intercept effect (*lme4::glmer*; Bates et al. 2015). This framework allowed us to estimate the B-positive frequency for a typical family (inverse-logit of the intercept) with its 95% confidence interval and the among-family variance.

### Population structure

Filtered biallelic SNPs called against J621C were used for principal-component analysis in PLINK 2.0 and model-based clustering in ADMIXTURE (Alexander et al. 2009; K = 1–7; K = 5 selected by the lowest cross-validation error, **Table S10**). Individuals were assigned to five genetic subgroups, re-estimated here rather than inherited from Lovell et al. (2021); the first three principal components (capturing the major north-south axis of structure) were retained as covariates for the expression analyses.

### Phylogenetic analysis

To reconstruct the evolutionary relationships within the Atlantic subpopulation, we first converted the filtered VCF file into a PHYLIP alignment format using the *vcf2phylip* tool (v2.0) (Ortiz 2019) with default parameters. This alignment was then used as input for phylogenetic inference using the coalescent-aware method CASTER (Zhang et al. 2025). The phylogenetic tree was inferred using the caster-site algorithm, employing 16 independent search rounds followed by 16 rounds of subsampling and exploration to ensure convergence on the global optimum. Branch support was estimated using 1000 replicates of local block bootstrapping. The inferred topology is unrooted, but is shown midpoint-rooted in **Fig. 3C** for visualization purposes. We chose CASTER for this analysis due to its demonstrated superior accuracy in handling incomplete lineage sorting and gene flow compared to traditional concatenation or other summary-coalescent methods like SVDQuartets (Zhang et al. 2025).

### Total RNA isolation and sample quality validation

To investigate the transcriptional activity and functional impact of the supernumerary B-Chromosome, leaf tissues were harvested from a core panel of 135 unique Atlantic switchgrass genotypes. This RNA-seq resource was generated as part of a companion study of expression variation in the same Atlantic panel (Izquierdo et al. 2026) and is re-analyzed here with respect to B-Chromosome carrier status. We collected two biological replicates per genotype to ensure reproducible profiles across leaf collections. Sampling was carefully timed to capture individuals at the identical pre-flowering developmental stage across both B-Chromosome carrier lines and non-carrier controls. Field collections were split across two separate dates, May 6, 2021, and June 24, 2021, and were strictly carried out between 10 AM and 12 PM to control for any confounding diurnal expression changes. For each sample, the basal leaf tissue was placed into a 15 mL tube and immediately flash-frozen in liquid nitrogen within 60 seconds of removal from the live plant. Frozen samples were transported on dry ice and stored at −80°C until RNA isolation as described by Izquierdo et al. (2026).

### RNA-seq data processing, differential expression, co-expression and GO analysis

RNA-seq was performed using NovaSeq S4 paired-end sequencing (2 x 151 bp) at the DOE Joint Genome Institute. Raw sequencing reads were processed, filtered, and checked for quality using fastp to trim adapter sequences and remove low-quality bases. The filtered high-quality reads from each sequencing library were then aligned to our newly generated chromosome-level Atlantic switchgrass reference genome (J621C v2.1) using the STAR aligner configured in 1-pass mode. Following alignment, raw fragments mapping to annotated genomic features were quantified using Subread-featureCounts to generate the primary gene expression count matrix.

Outliers among biological replicates were identified and excluded based on their Euclidean distance from the cluster center and their Pearson correlation with the other replicates (libraries with *r* < 0.85 were excluded). Only libraries that passed quality-control criteria and outlier filtering were retained for downstream analyses. The filtered raw counts were normalized into Transcripts Per Million (TPM) and Fragments Per Kilobase of transcript per million mapped reads (FPKM) values to scale for sequencing depth and gene length. For differential expression profiling and gene co-expression network analysis, the raw count matrix was systematically transformed via Variance Stabilizing Transformation (VST) and Relative Log Expression (RLE) protocols implemented directly within the DESeq2 package (Love et al. 2014) to stabilize the variance across the wide range of expression levels. This normalized dataset was used for all subsequent transcriptomic evaluations of the B-Chromosome’s functional impacts. Differential expression between carriers and non-carriers was tested separately within the SP and NP groups with DESeq2 using the design ∼ PC1 + PC2 + PC3 + B status, after retaining genes with ≥10 reads in ≥3 libraries; each B+ genotype was matched 1:1, without replacement, to the nearest B− genotype of the same group in PC1–PC3 space, and all replicate libraries of the selected genotypes were kept. Log2 fold changes were shrunk with *apeglm* (Zhu et al. 2019), B-linked genes were excluded, and genes with BH-adjusted P < 0.05 and |log2 fold change| > 1 were called differentially expressed. For co-expression, variance-stabilized counts (design ∼ group + PC1 + PC2 + PC3 + B status) of B-linked genes with ≥10 reads in ≥5 libraries were analyzed with WGCNA (Langfelder and Horvath 2008; soft-threshold power 6, unsigned network, minimum module size 10, merge height 0.25), and module eigengenes were correlated with B status and genetic group (Pearson). Genes not assigned to any module do not share an expression pattern and were therefore not summarized by an eigengene; their association with B status and genetic group was tested gene by gene (Pearson correlation, Benjamini-Hochberg correction across genes and traits). GO enrichment used clusterProfiler (Wu et al. 2021; *enricher* and *compareCluster*) with the GO-annotated J621C genes as the universe, gene sets of 3-500 genes, and terms retained at BH-adjusted *P* < 0.05 and *q* < 0.20 (**Tables S16-S19**).

## Data availability

All data generated in this study will be deposited in public repositories before publication. The J621C and J586A genome assemblies and annotations will be released through DOE-JGI Phytozome and NCBI GenBank, and the PacBio HiFi, Illumina and Omni-C reads through the NCBI Sequence Read Archive (SRA); accession numbers will be provided before publication. The resequencing data re-analyzed here are available from NCBI SRA under the identifiers listed by Lovell et al. (2021) and the RNA-seq data were described by Izquierdo et al. (2026). All other data supporting the findings are provided in the Supplementary Tables, and the analysis code will be deposited in a public repository before publication.

## Acknowledgments

This work was supported by the U.S. Department of Energy (DOE), Office of Science, Biological and Environmental Research, Genomic Science Program under Award Numbers DE-SC0021126 and DE-SC0026058, as well as the Great Lakes Bioenergy Research Center under Award Number DE-SC0018409. The work (proposal: 10.46936/10.25585/60000507) conducted by the U.S. DOE Joint Genome Institute (https://ror.org/04xm1d337), a DOE Office of Science User Facility, is supported by the Office of Science of the U.S. DOE operated under Contract No. DE-AC02-05CH11231. Additional support was provided by the National Science Foundation Long-term Ecological Research Program (DEB-2224712) at the Kellogg Biological Station and by Michigan State University AgBioResearch. JTL is supported by the Center for Bioenergy Innovation (CBI), which is supported by the U.S. Department of Energy, Office of Science, Biological and Environmental Research Program under Contract Number ERKP886. The authors acknowledge the Texas Advanced Computing Center (TACC) at The University of Texas at Austin for providing computational resources that contributed to the research results reported within this paper (http://www.tacc.utexas.edu).

## Author Contributions

A.F.E., A.L.H., J.S., and T.E.J. designed research; A.F.E., J.E.B., U.B.S., L.B., J. Guo, and J.T. performed research; J.T.L. and J. Guo contributed new analytic tools; A.F.E., L.Z., A.L.H., A.S., J.J., and S.Z. analyzed data; T.E.J., D.B.L., J. Grimwood, and J.S. supervised research; K.B. administered the project; and A.F.E. and T.E.J. wrote the paper with input from all authors.

## Supplementary figures

**Fig. S1.**
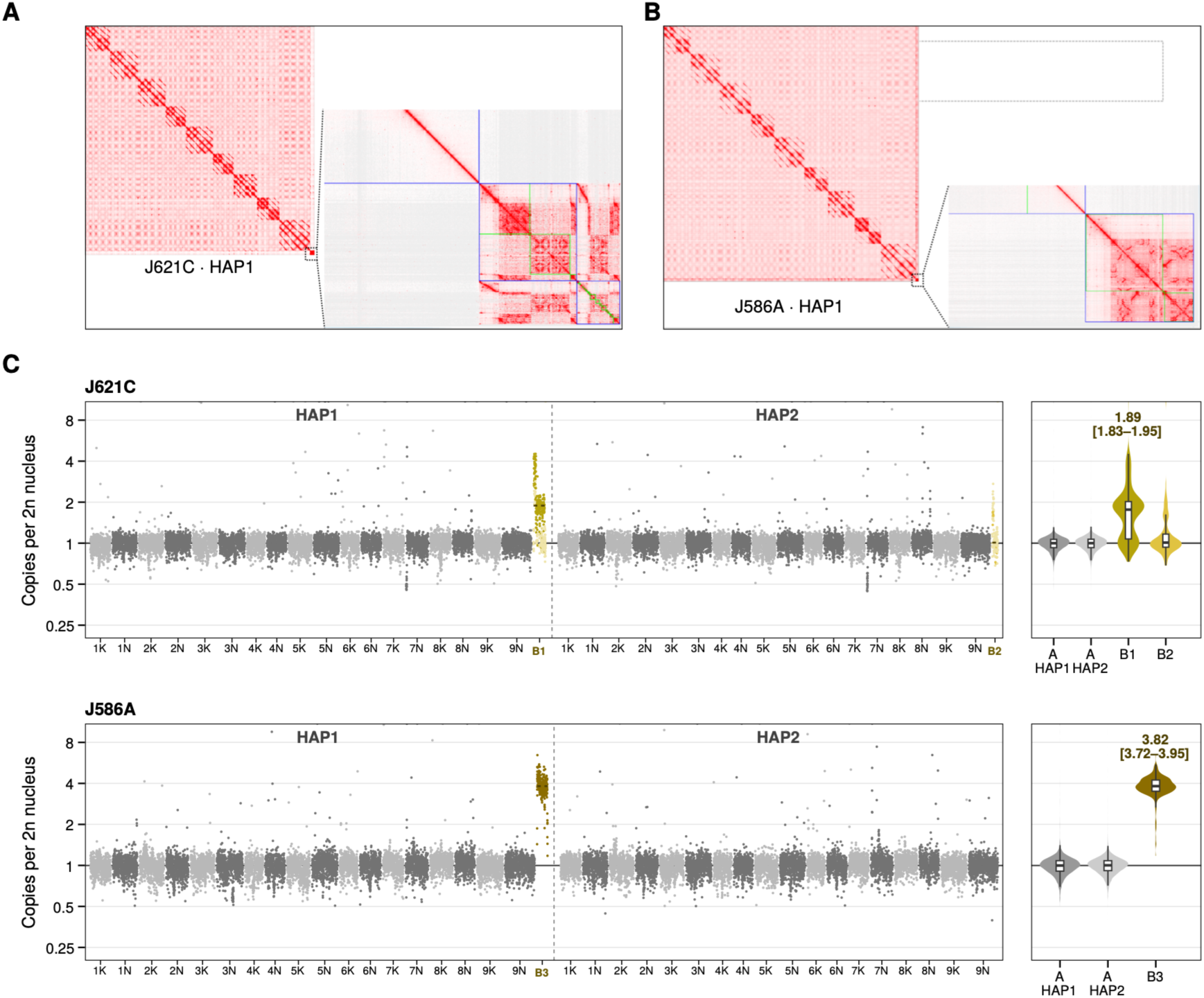
Validation of the B-Chromosome assemblies by Hi-C contact maps and HiFi read depth. (**A** and **B**) Hi-C contact maps of J621C HAP1 (**A**) and J586A HAP1 (**B**). Full genome contact maps (Juicebox) with Chr19 shown enlarged. Each B forms a distinct contact domain relative to the 18 A-Chromosomes (K and N subgenomes), supporting its assembly as an independent chromosome. (**C**) HiFi read depth of J621C and J586A in 100-kb windows on the combined reference of each genotype (HAP1 + HAP2 + all assembled B copies), expressed as copies per diploid nucleus (window depth divided by the median single-copy depth of the A-Chromosomes). **Left**, all chromosomes of HAP1 and HAP2, with each B copy placed next to the haplotype in which it was assembled (J621C: B1 after the HAP1 and B2 after the HAP2 A-Chromosomes; J586A: B3 after the HAP1 A-Chromosomes). A-Chromosomes are shown in alternating grey shades and B copies in gold; pale gold marks J621C windows shared by B1 and B2, whose reads are divided between the two copies; horizontal dashed lines mark the median B copy number, and the vertical dashed line separates HAP1 from HAP2. **Right**, distribution of window copy number per sequence class (violin and box plots, on the same y axis); numbers give the median B copy number with its 95% confidence interval. Values are in **Table S2**.

**Fig. S2.**
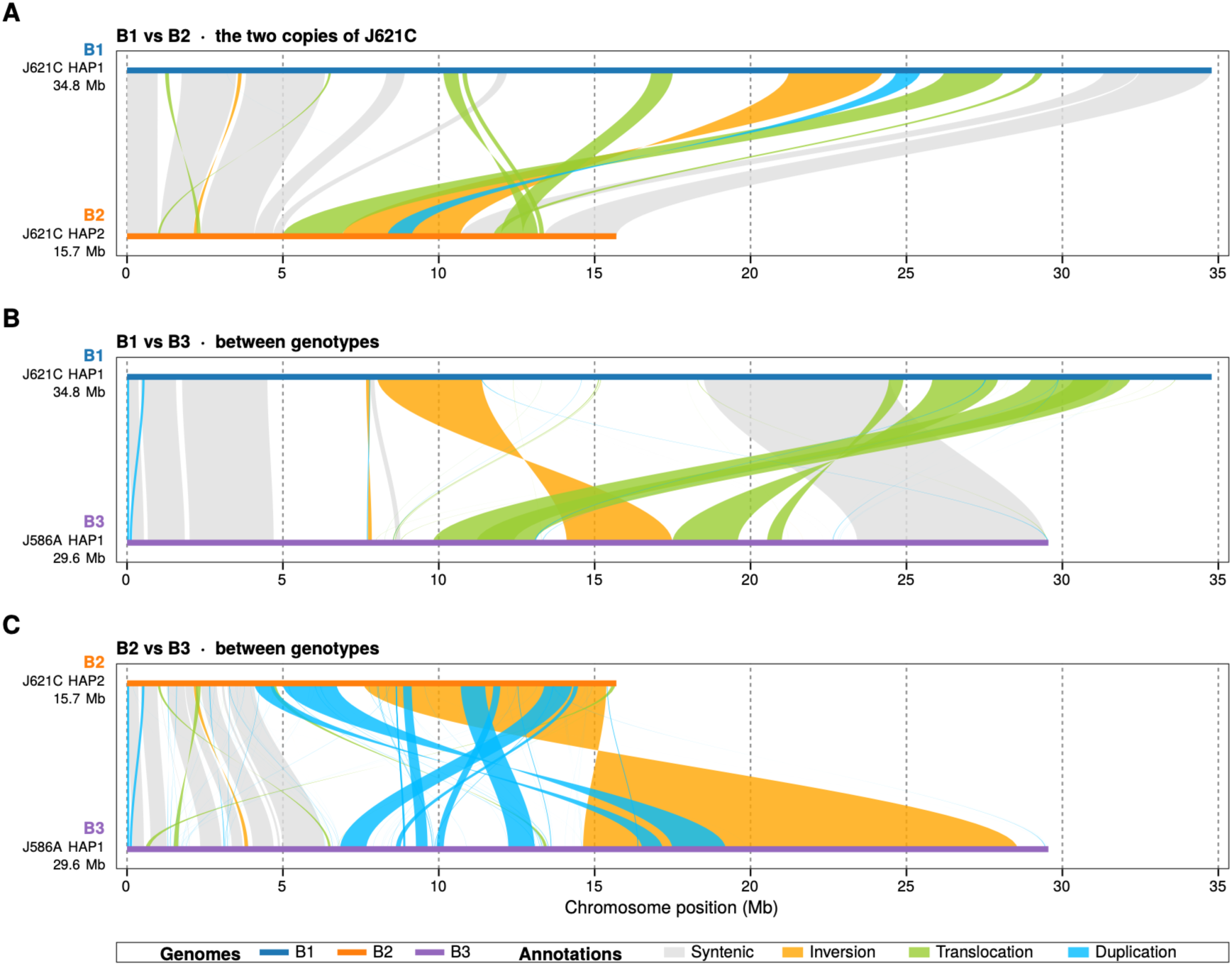
Structural comparison of the three assembled B-Chromosome copies (B1, B2, and B3). Whole-sequence alignments (minimap2) annotated with SyRI for (**A**) B1 (J621C HAP1, 34.8 Mb) versus B2 (J621C HAP2, 15.7 Mb), (**B**) B1 versus B3 (J586A HAP1, 29.6 Mb) and (**C**) B2 versus B3. Each copy is a colored line on a shared position axis (Mb; the first-named copy on top), and ribbons join the aligned blocks, colored by SyRI annotation (syntenic, inversion, translocation or duplication; legend below **C**); crossed ribbons are inverted blocks. Aligned fractions, SNP densities and k-mer distances are given in **Table S3**. Complementary sequence dot plots appear in **Fig. S3**.

**Fig. S3.**
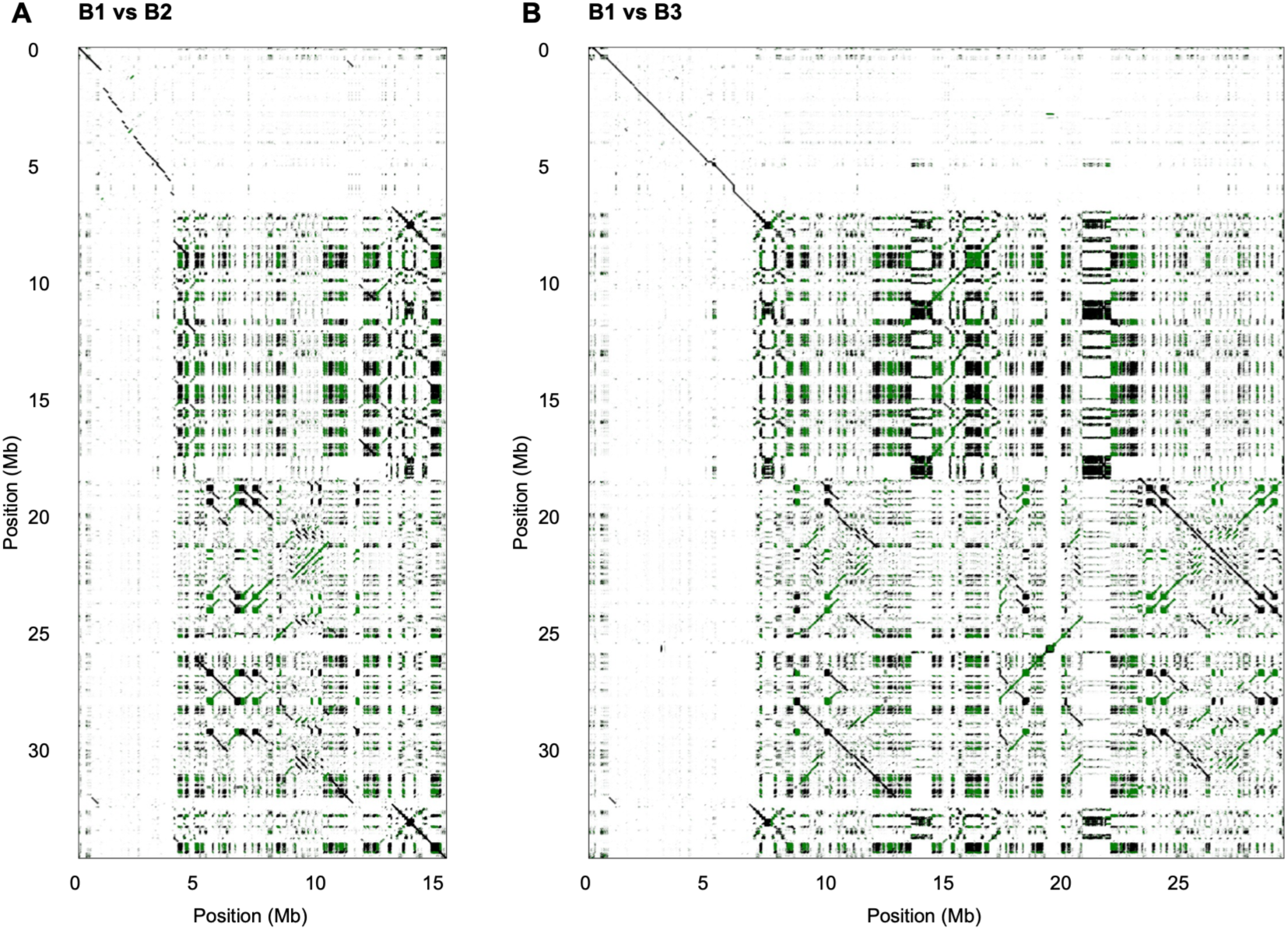
Pairwise FlexiDot comparisons of B1 with (A) B2 and (B) B3. B1 (J621C HAP1) occupies the vertical axis in both panels; B2 (J621C HAP2) and B3 (J586A HAP1) occupy the horizontal axes. Position labels are in megabases. These comparisons complement the classified rearrangements in **Fig. S2**.

**Fig. S4.**
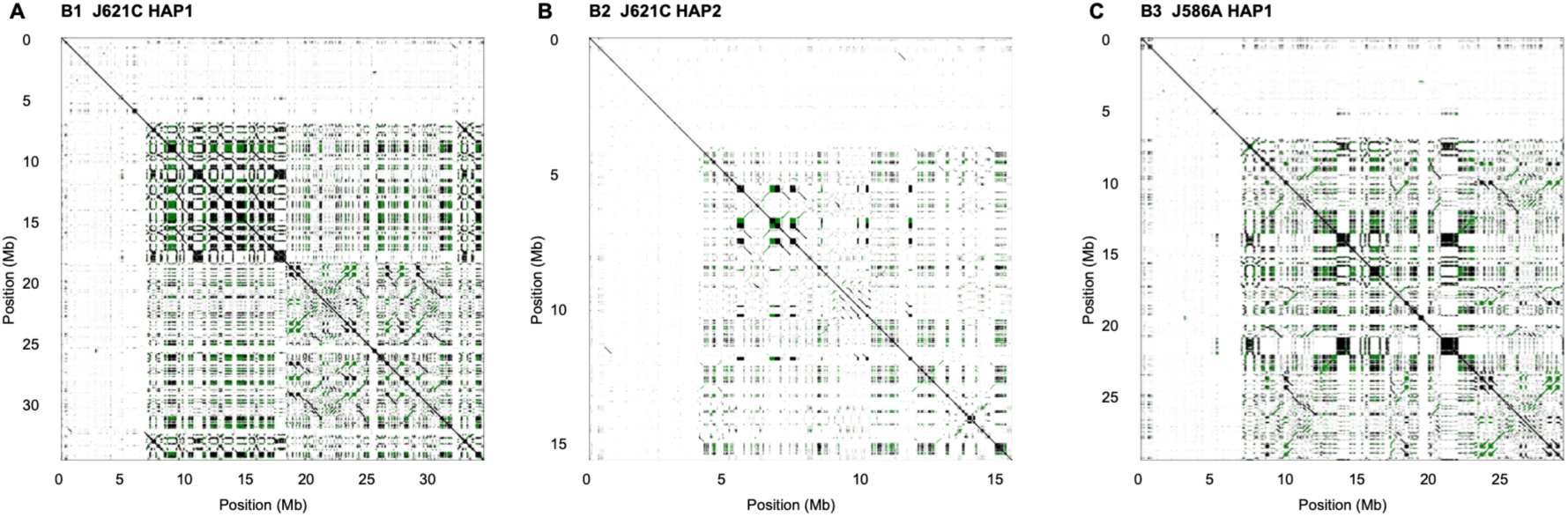
FlexiDot self dot plots of the three assembled B-Chromosome copies. (**A**) B1 (J621C HAP1), (**B**) B2 (J621C HAP2) and (**C**) B3 (J586A HAP1), each plotted against itself with a *k*-mer of 81 bp.

**Fig. S5.**
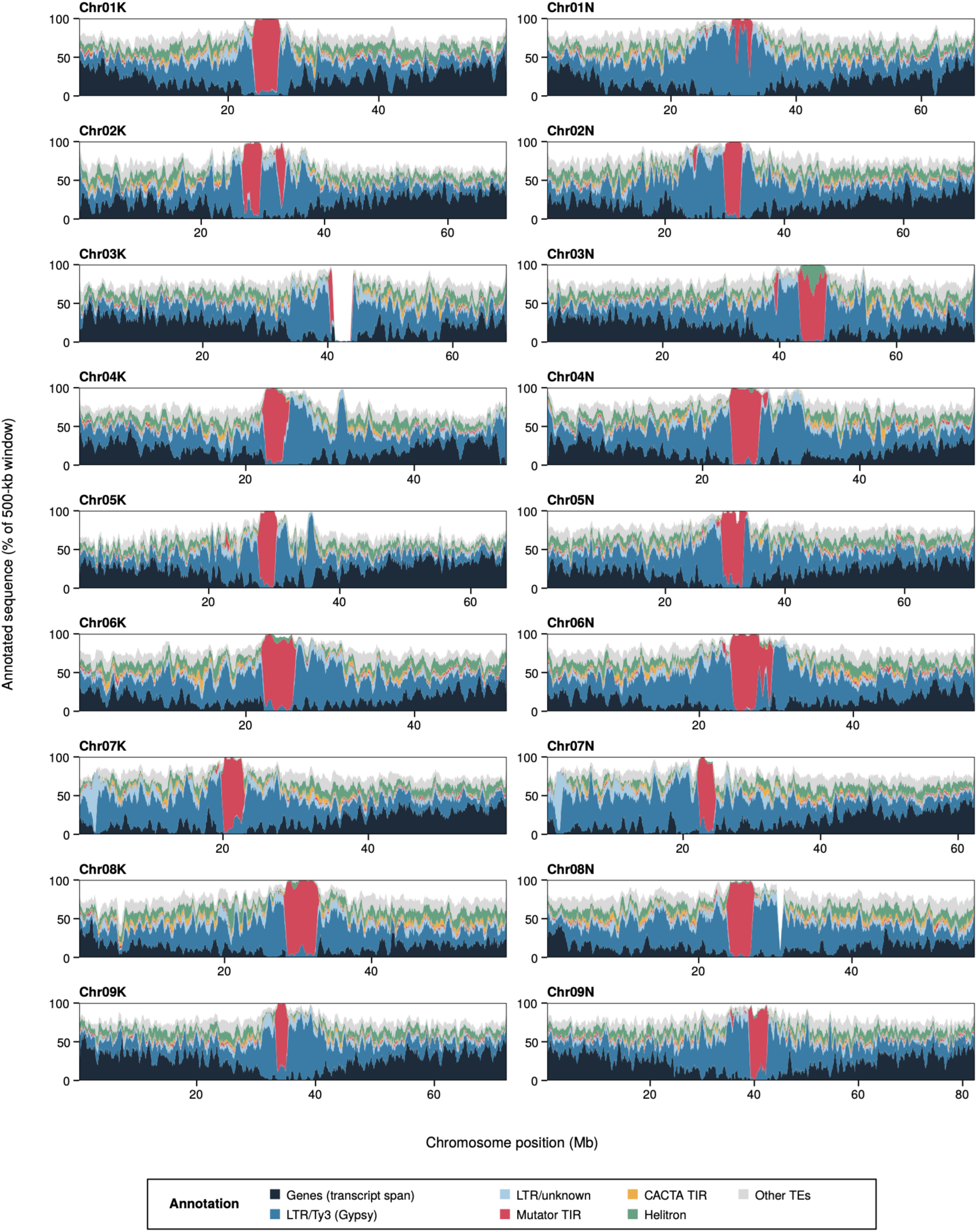
Repeat and gene annotation landscapes of all 18 J621C A-Chromosomes. Each panel shows the percentage of each 500-kb window (50-kb steps) annotated as each class, computed as in Fig. 2A; colors follow the annotation key of Fig. 2A**-B**. The x-axis of each panel spans that chromosome (Mb).

**Fig. S6.**
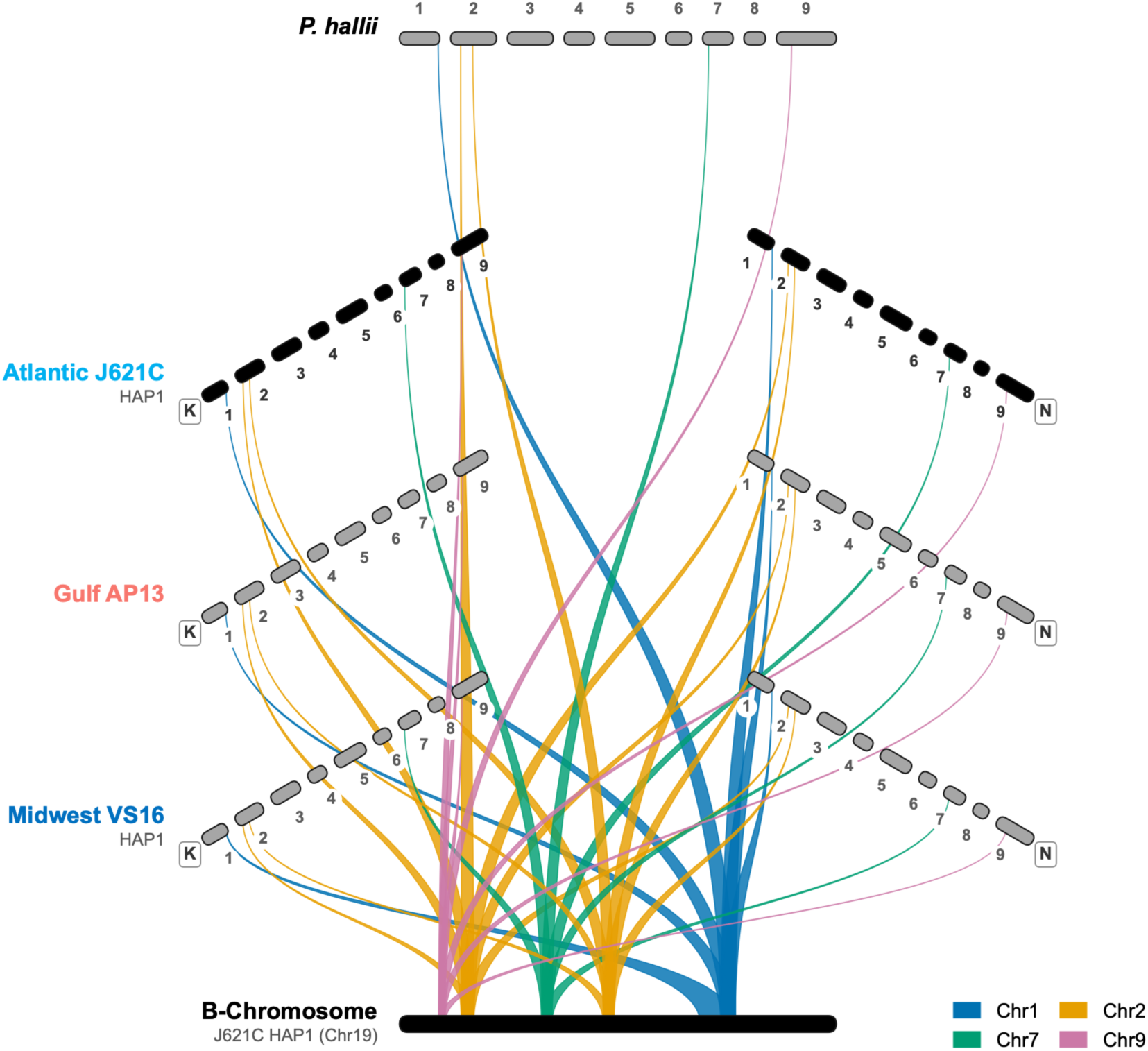
Gene synteny analysis detailing the ancestral A-Chromosome origins of the switchgrass B-Chromosome. The plot displays the sequence relationships of the supernumerary element by mapping gene synteny between the B-Chromosome (J621C HAP1; bottom) and the A-Chromosomes across multiple switchgrass reference genomes (J621C, AP13, VS16) and the closely related outgroup *Panicum hallii* (top). The distinct colored linkages demonstrate that rather than originating from a single autosome, the B-Chromosome is a highly rearranged chimera. It is composed of syntenic blocks derived from several ancestral autosomes, principally Chromosomes 1, 2, 7, and 9.

**Fig. S7.**
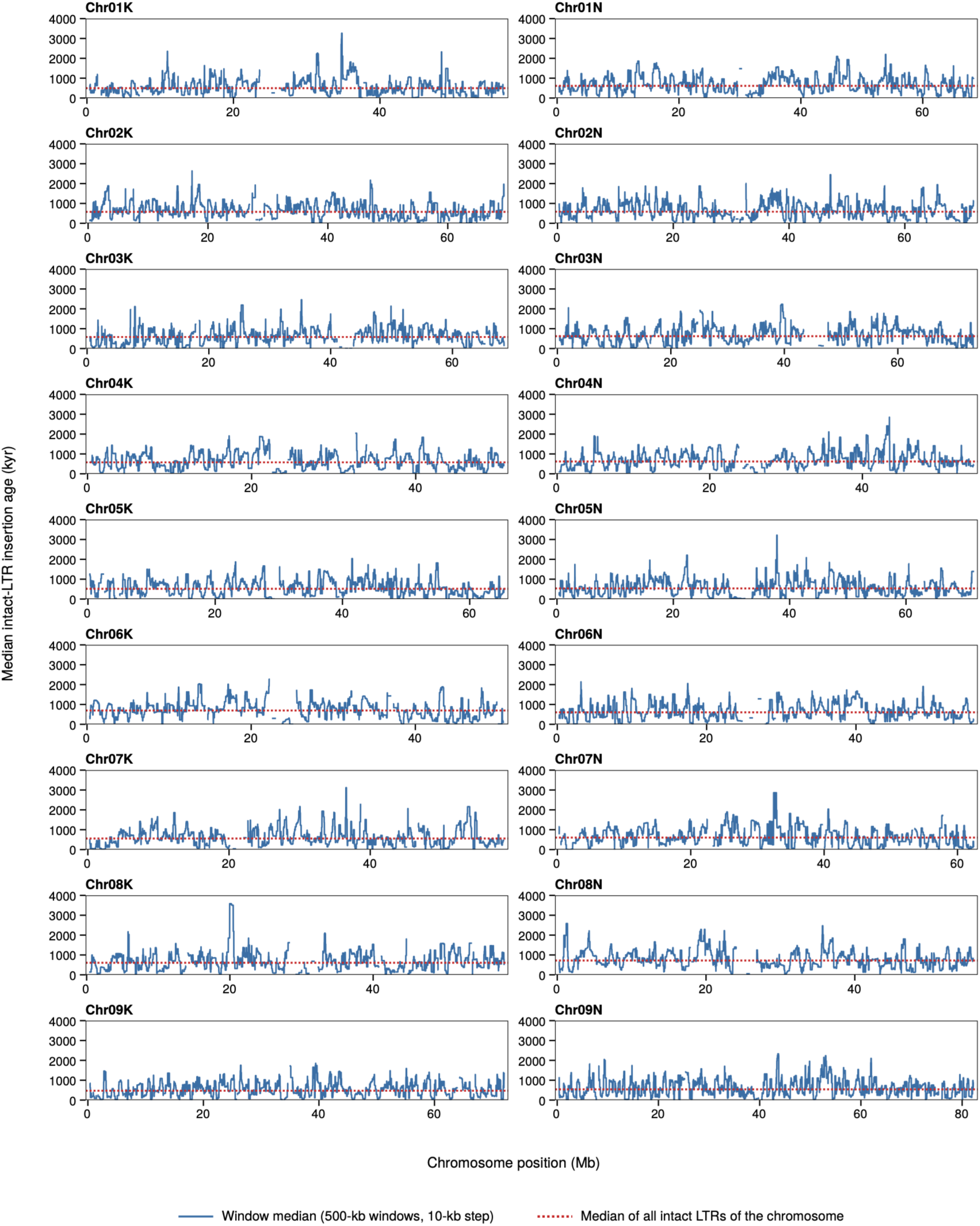
Intact LTR insertion-age profiles for all 18 J621C A-Chromosomes. Age estimates use 500 kb windows stepped by 10 kb and are shown in thousands of years (kyr). Red dotted line represents overall median age of intact LTRs per chromosome. Horizontal positions are in megabases; the x-axis of each panel spans that chromosome (layout as in **Fig. S5**).

**Fig. S8.**
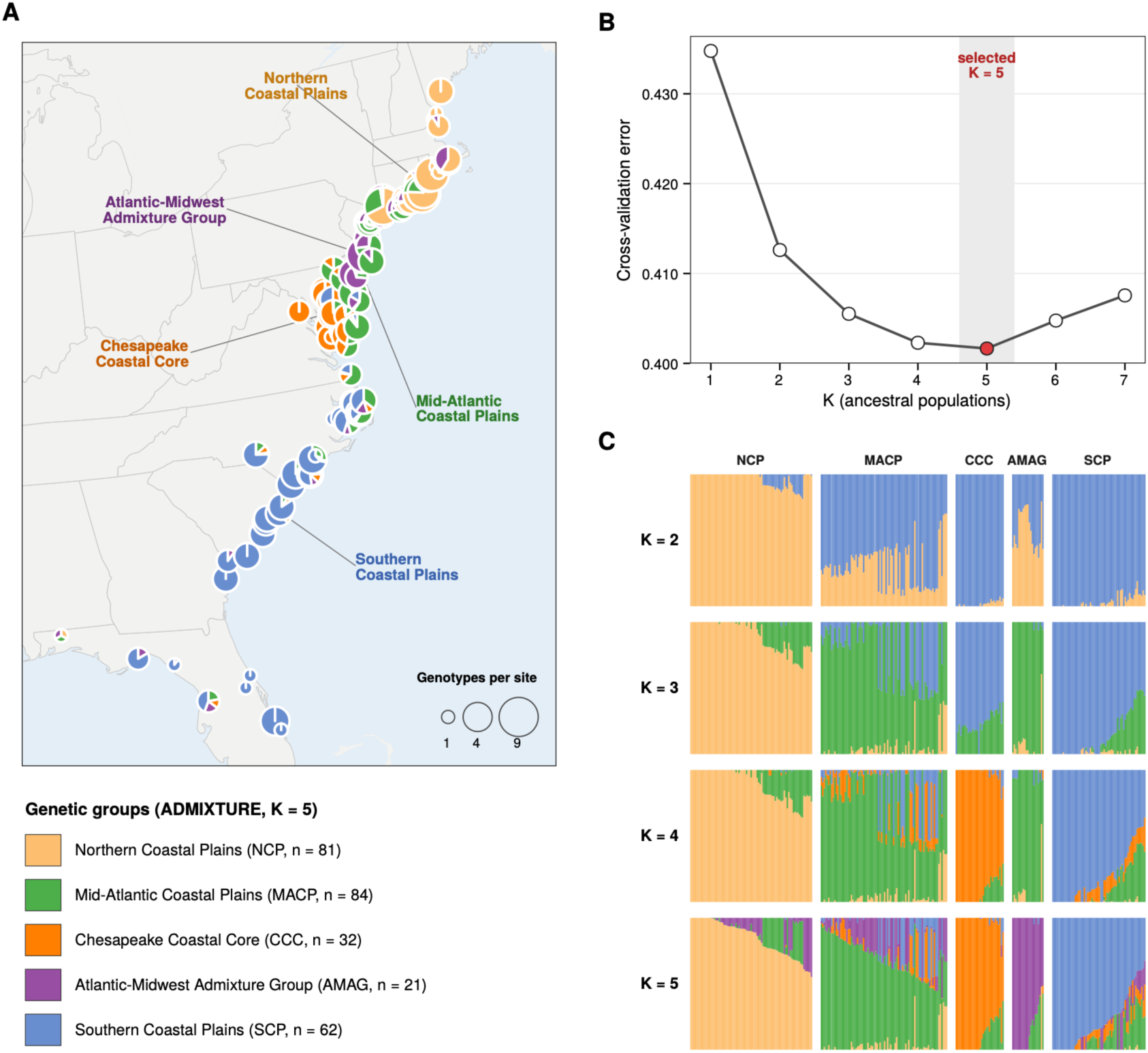
Admixture analysis of the Atlantic switchgrass subpopulation. (**A**) Geographic distribution of the sampled *Panicum virgatum* individuals along the North American Atlantic coast. Pie charts at each sampling location indicate the average ancestry proportions (K = 5) of the genotypes sampled at that location; the five genetic groups are labeled. (**B**) Mean cross-validation (CV) error across 10-fold cross-validation replicates for increasing numbers of genetic clusters (K = 1 to 7). The optimal model complexity is identified at K = 5, where the CV error reaches its minimum. (**C**) ADMIXTURE bar plots displaying individual ancestry proportions for K = 2 through K = 5. Each vertical bar represents a single individual, segmented and colored according to the estimated proportion of ancestry assigned to each cluster. Individuals are grouped along the x-axis by their primary genetic cluster assignment at the optimal K = 5 model and are shown in the same order in every row; each ancestry component is colored after the K = 5 group that carries most of it.

**Fig. S9.**
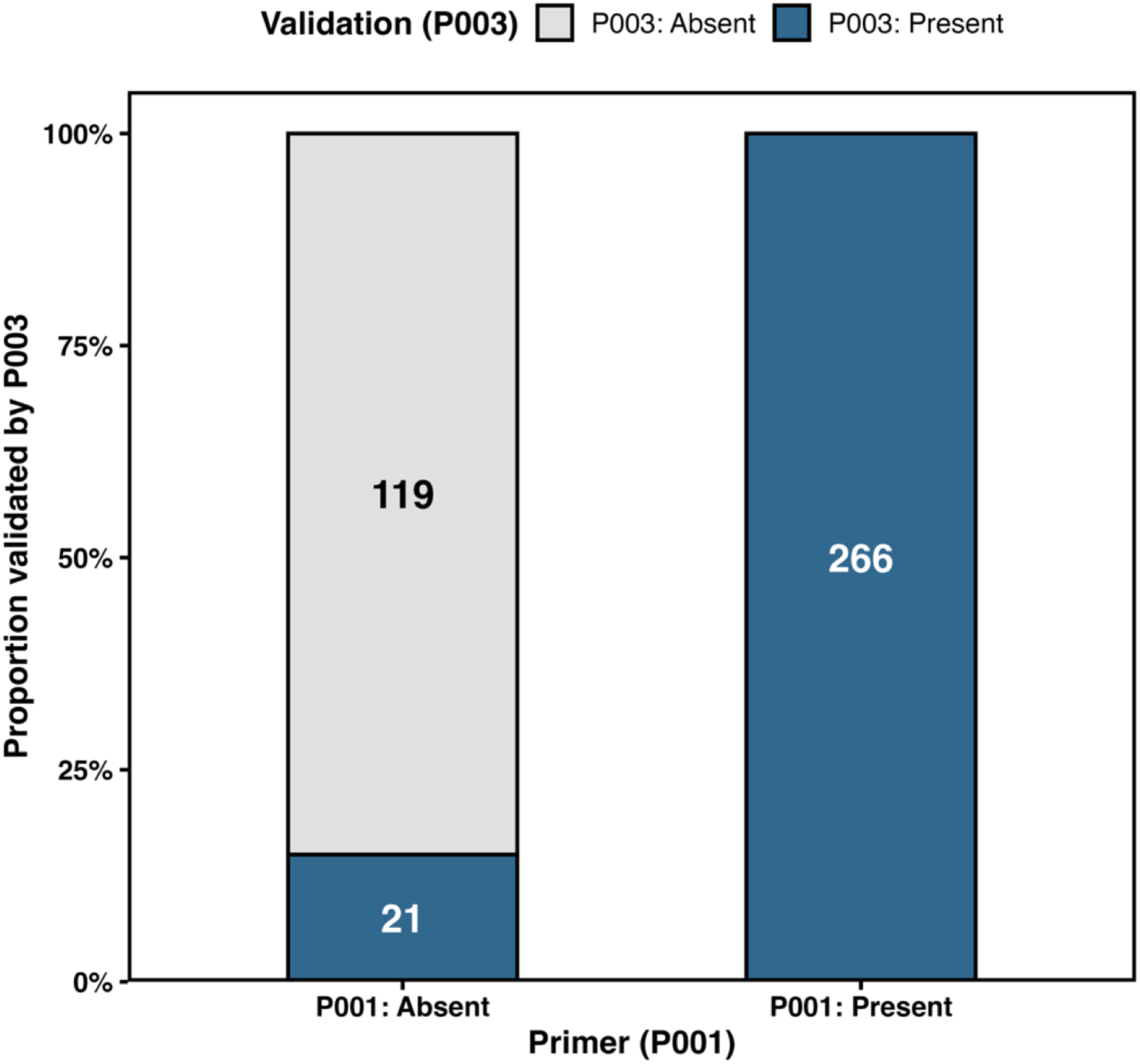
Genotyping validation of B-Chromosome presence. The stacked bar chart displays the proportion and absolute counts of individuals scored as carrying the B-Chromosome (blue) or lacking it (grey) by the validation primer (P003), conditional on their initial P001 screening status. The strong statistical agreement between the two markers (Cohen’s Kappa = 0.881, *p* < 0.001, N = 406) supports agreement between the two assays (per-progeny calls in **Table S14**).

## Supplementary tables

**Table S1.** Sequencing, assembly and quality statistics of the J621C and J586A haplotype-resolved genome assemblies.

**Table S2.** PacBio HiFi read depth and copy number of the assembled B-Chromosome copies.

**Table S3.** Sequence and structural comparison of the three assembled B-Chromosome copies (B1, B2, B3).

**Table S4.** Protein-coding genes annotated on the J621C B-Chromosome (B1; Chr19 of J621C HAP1 v2.1).

**Table S5.** Repeat and gene composition of the A-Chromosomes and the B-Chromosome of J621C HAP1.

**Table S6.** Candidate 166-bp centromeric tandem repeat and its intervals on the B-Chromosome (TRASH).

**Table S7.** Autosome and subgenome of origin of B-Chromosome single-copy sequence (*k*-mer painting).

**Table S8.** Insertion ages of intact LTR retrotransposons on each J621C HAP1 chromosome.

**Table S9.** Atlantic resequencing panel: sampling locations, genetic structure and B-Chromosome status (280 natural-collection genotypes).

**Table S10.** ADMIXTURE cross-validation error and PCA eigenvalues for the Atlantic SNP set.

**Table S11.** Genome-wide B-specific *k*-mer screen of the switchgrass diversity panel of Lovell et al. (2021) (696 of 732 genotypes screened).

**Table S12.** Frequency of B-positive genotypes by Atlantic genetic group and by switchgrass subpopulation.

**Table S13.** B-Chromosome-specific primer pairs used to genotype F1 progeny.

**Table S14.** PCR genotypes of 406 open-pollinated F1 progeny from 23 B-positive maternal lines.

**Table S15.** B-positive frequency among open-pollinated progeny of 23 B-positive maternal lines: family summaries, marker agreement and GLMM results.

**Table S16.** RNA-seq libraries, carrier status and inclusion in the expression analyses.

**Table S17.** Co-expression module of B-linked genes (WGCNA) and gene-by-gene tests of the unassigned genes.

**Table S18.** A-Chromosome genes differentially expressed between B-positive and B-negative genotypes.

**Table S19.** Gene Ontology enrichment of the background-specific A-Chromosome DEG sets.

